# Quantifying the intricacies of biological pattern formation: A new perspective through complexity measures

**DOI:** 10.1101/2024.11.03.621719

**Authors:** Jurgen Riedel, Chris P. Barnes

## Abstract

Nonlinear pattern-forming systems can generate spatial structure through many mechanisms, but comparing the patterns that are actually realized after the dynamics unfold remains difficult. Linear stability analysis identifies when spatial modes can grow and which length scales are favored near onset, but it does not directly measure the state diversity, structural organization, or dynamical route of the resulting patterns. Here we present a comparative operational framework, mechanism-agnostic under a fixed measurement protocol, based on two coordinates: the Diversity of Number of States (DNOS), an occupied-state measure of effective alphabet breadth, and the Diversity of Pattern Complexity (DPC), a lossless-compression measure of structural complexity. We apply the framework to a linear reaction–diffusion benchmark, the Gray–Scott and FitzHugh–Nagumo models, toggle-switch gene-regulatory systems with and without self-activation, and a Notch–Delta–EGF model of Drosophila neurogenesis. Across these systems, DNOS–DPC landscapes identify realized pattern regimes not determined by linear onset analysis alone. Robustness checks against bin count, entropy-based DNOS diagnostics, compression settings, image noise, clustering method, and graph, spectral, and wavelet descriptors support the stability of the comparative summaries under the stated protocol. In the Drosophila model, end-point, field-resolved, multichannel, and time-resolved analyses distinguish pathway-level Notch/Delta signaling collapse from partial downstream proneural-output persistence, and suggest an EGF-dependent entry/progression bias into organized proneural-wave states. The framework provides reproducible coordinates for comparing realized pattern diversity, structural complexity, and developmental trajectories across nonlinear reaction–diffusion and regulatory systems.

## 1 Introduction

A central question in nonlinear pattern formation is how local interactions, feedback, transport, and instability generate reproducible spatial organization. Developmental biology provides many prominent examples of this problem, because initially similar cells and tissues can organize into animal markings, cell-state boundaries, and differentiating tissue patterns. Reaction–diffusion and gene-regulatory models have played a major role in addressing this question since Turing’s foundational work [1] and the subsequent formulation by Gierer and Meinhardt [2]. These models show how local interactions, feedback, and transport can generate spatial organization across a wide range of biological settings [3–8]. However, a persistent difficulty is that meaningful comparison across models remains hard: two systems may both form spatial structure, yet differ substantially in how many states they realize, how ordered or irregular those states are, and how strongly those outcomes depend on parameters or network architecture.

Standard analysis of reaction–diffusion systems relies on linear stability theory and bifurcation methods [9]. These approaches identify when a homogeneous state loses stability and which wavelengths are favored near onset, but they do not directly describe the realized patterns that persist at long times. Recent work shows that instability alone is not a reliable indicator of durable pattern formation, especially in systems with multistability, strong nonlinearities, stochastic effects, or far-from-onset dynamics [10–16]. The key question is therefore not only whether a system can become unstable, but what kind of spatial organization it actually realizes once the full nonlinear dynamics unfold.

This creates a comparison problem that is broader than pattern detection. Nonlinear developmental and regulatory systems must be compared after their dynamics have unfolded, because the same linear onset class can lead to different realized spatial states, and visually similar endpoints can arise through different temporal routes. In biological applications, this distinction can be especially important: a mutation may leave a downstream output field partially organized while disrupting an upstream signaling channel. A useful theoretical descriptor should therefore compare final pattern organization, field-specific contributions, and dynamical trajectories within one common coordinate system. This is the motivation for treating DNOS and DPC not only as endpoint descriptors, but also as time-resolved coordinates for realized pattern formation.

Other limitations arise in systems with strong nonlinearities, high-dimensional interactions, or far-from-equilibrium dynamics. Amplitude-equation approaches may fail away from bifurcation points, and deterministic analyses frequently neglect stochastic effects that can influence pattern-forming systems [14–16]. These challenges motivate complementary ways to characterize patterns that do not require detailed analytical tractability.

Complexity-based methods offer a general framework for describing patterns in terms of their structural detail and state diversity, without relying on predefined geometric categories. Such approaches have been used to compare patterns across models and to quantify nonlinear interactions in biological networks [17–19]. They are applicable to large parameter spaces and can be used to assess the organization of systems with intricate feedbacks or multistability, including gene regulatory networks [20, 21].

Many previous studies of biological patterning have used geometric or morphology-based descriptors tailored to specific systems, such as fish, reptile, or mammalian pigmentation [22–25]. Theoretical work has also examined pattern selection and stability through weakly nonlinear analysis, focusing on transitions between stripes, spots, and labyrinthine motifs and the influence of domain shape [26, 27]. These methods provide mechanistic insight, but they typically require model-specific derivations and are not easily generalized across distinct systems.

In this work we take a complementary approach. Rather than extracting geometric features or relying on system-specific measurements, we analyze the realized patterns directly through their intensity distributions and information content. This produces a mechanism-agnostic description under a fixed measurement protocol, allowing realized patterns from different reaction–diffusion systems and gene-regulatory networks to be compared on common operational axes. The goal is not to assign an absolute or universal complexity value to a pattern, but to provide a reproducible coordinate system for comparing realized outcomes within and across model classes. Our aim is to characterize how such systems populate intensity space, how structurally compressible their spatial configurations are, and how those quantities change across parameters, regulatory architecture, and developmental perturbation.

Two questions guide this study. First, given a system that becomes unstable and develops spatial structure, what is the character of the pattern that emerges? Second, can we summarize the complexity of these patterns in a way that allows comparison across different models and parameter sets? To address these questions, Section 2.2 introduces the Diversity of Number of States (DNOS) and the Diversity of Pattern Complexity (DPC) as the primary operational measures used throughout this study to characterize realized patterns. The Uniformity Measure Entropy (UME) is used more narrowly as an auxiliary normalized descriptor of uniformity that helps interpret or refine these summaries where appropriate, rather than as a coequal headline measure.

**Structure of the paper.** After motivating the problem in the Introduction, we first establish the linear reaction–diffusion benchmark and the complexity measures used throughout the paper. Section 2.1 presents the spectral analysis of the linearized reaction–diffusion reference system, and Section 4.3 defines the complexity measures DNOS, DPC, and UME. We then apply these measures across the model classes studied here: Sections 2.3, 2.4, and 2.5 analyze the linearized reaction–diffusion, Gray–Scott, and FitzHugh–Nagumo systems; Sections 2.6.1 and 2.6.2 examine toggle–switch gene-regulatory models with and without self–activation; Figure 7 compares all systems in a common complexity space; and Section 2.8 analyzes the Notch–Delta–epidermal growth factor signaling model of *Drosophila* neurogenesis using endpoint, field-resolved, multichannel, and time-resolved DNOS–DPC summaries. Section 3 discusses the implications, limitations, and possible extensions of the framework. Technical details are collected in Section 4.1 and Section 4.3, which describe the numerical methods, spectral benchmark, formal definitions of the complexity measures, auxiliary descriptors, clustering procedures, and finite-difference implementation.

## 2 Results

We organize the Results around one central question: can DNOS and DPC provide a common comparative description of realized biological patterns across mechanistic classes, beyond what linear onset analysis alone can show?

### 2.1 Stability analysis of the linearized reaction–diffusion system

Turing’s original analysis showed how reaction–diffusion systems can break spatial homogeneity through a diffusion-driven instability of a homogeneous steady state [1]. Here we use a linear two-species Turing system as an analytically tractable benchmark for spectral stability and length-scale analysis. Its role is to provide a reference for where spatial structure is expected to emerge, distinct from the time-evolved simulations used later to quantify realized patterns through DNOS and DPC.

A generic two-species reaction–diffusion system is

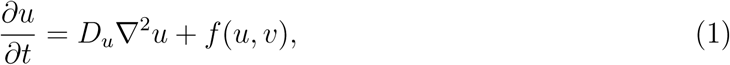

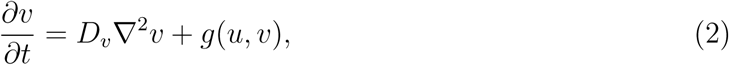

where *u* and *v* denote the concentrations of two interacting species, *D_u_* and *D_v_* are their diffusion coefficients, and *f, g* encode the local reaction kinetics. Let (*h, k*) be a homogeneous steady state, so that *f* (*h, k*) = *g*(*h, k*) = 0. Linearizing the kinetics around (*h, k*) yields

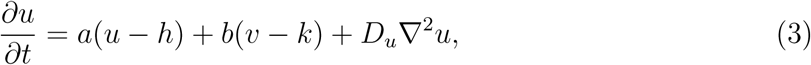

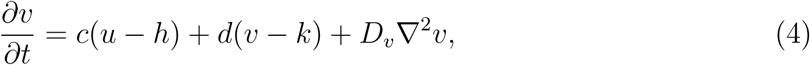

where *a* = *f_u_*(*h, k*), *b* = *f_v_*(*h, k*), *c* = *g_u_*(*h, k*), and *d* = *g_v_*(*h, k*) are the entries of the Jacobian of the reaction terms. Thus *a* and *d* describe self-feedback of *u* and *v*, while *b* and *c* are the cross-interaction coefficients.

To analyze linear stability we consider perturbations of the form (*u* −*h, v* −*k*) ∝ exp(*λt* + *i***k** · **x**), which leads to the dispersion relation

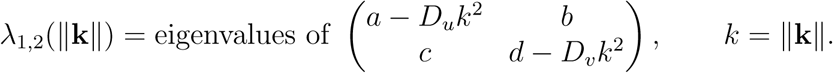

For each parameter set (*a, b, c, d, D_u_, D_v_*) and wavenumber *k* we compute the two eigenvalues *λ*_1_(*k*) and *λ*_2_(*k*) numerically. The homogeneous steady state is linearly stable if ℜ*λ*_1_(*k*) *<* 0 and ℜ*λ*_2_(*k*) *<* 0 for all *k*. A diffusion-driven (Turing) instability occurs when the homogeneous state is stable at *k* = 0 but ℜ*λ_i_*(*k*) *>* 0 for some finite *k >* 0. In a standard two-species system with ordinary diffusion, this finite-wavenumber instability is the relevant linear signature of Turing pattern formation. Accordingly, we restrict the analysis to quantities that are directly supported by the dispersion relation, rather than assigning nonlinear bifurcation labels such as pitchfork or transcritical classes.

The diffusion coefficients *D_u_* and *D_v_* are varied on a logarithmic grid in the range 10^−5^ to 10^−3^, while the reaction parameters *a, b, c, d* are sampled uniformly from [−2, 2]. For each parameter set we evaluate the dispersion relation on a grid of wavenumbers *k* ∈ [0, 100]. From the resulting eigenvalue spectra we derive, for each (*D_u_, D_v_*), four summary quantities: (i) the fraction of sampled Jacobians that are stable at *k* = 0, (ii) the fraction that are Turing-unstable in the numerical dispersion scan, that is, stable at *k* = 0 but unstable for some finite *k >* 0, (iii) the mean maximum growth rate ⟨max*_k>_*_0_ max*_i_* ℜ*λ_i_*(*k*)⟩ over the Turing-unstable cases, and (iv) the mean dominant wavelength ⟨2*π/k*_max_⟩, where *k*_max_ is the wavenumber with maximal growth rate for a given unstable sample. In addition, we separately evaluate the classical analytic Turing conditions as a validation of the numerical dispersion scan, but we do not use those validation maps as the main figure in the manuscript. The corresponding validation plots, including the analytic Turing-unstable fraction, the absolute difference between analytic and numerical classifications, and the fraction stable for all wavenumbers, are shown in Fig. S1 of the supplementary material.

Figure 1 summarizes these four spectral quantities over the (*D_u_, D_v_*) plane. Panel (A) shows the fraction of sampled Jacobians whose homogeneous steady state is stable at *k* = 0. Panel (B) shows the fraction that undergo a diffusion-driven instability at some finite wavenumber in the numerical dispersion scan. Panel (C) shows the mean maximum growth rate across the Turing-unstable cases. Panel (D) shows the corresponding mean dominant wavelength. This linear spectral analysis therefore identifies where diffusion-driven instabilities are expected, how strongly unstable modes grow, and which characteristic wavelengths are selected at onset.

**Figure 1:**
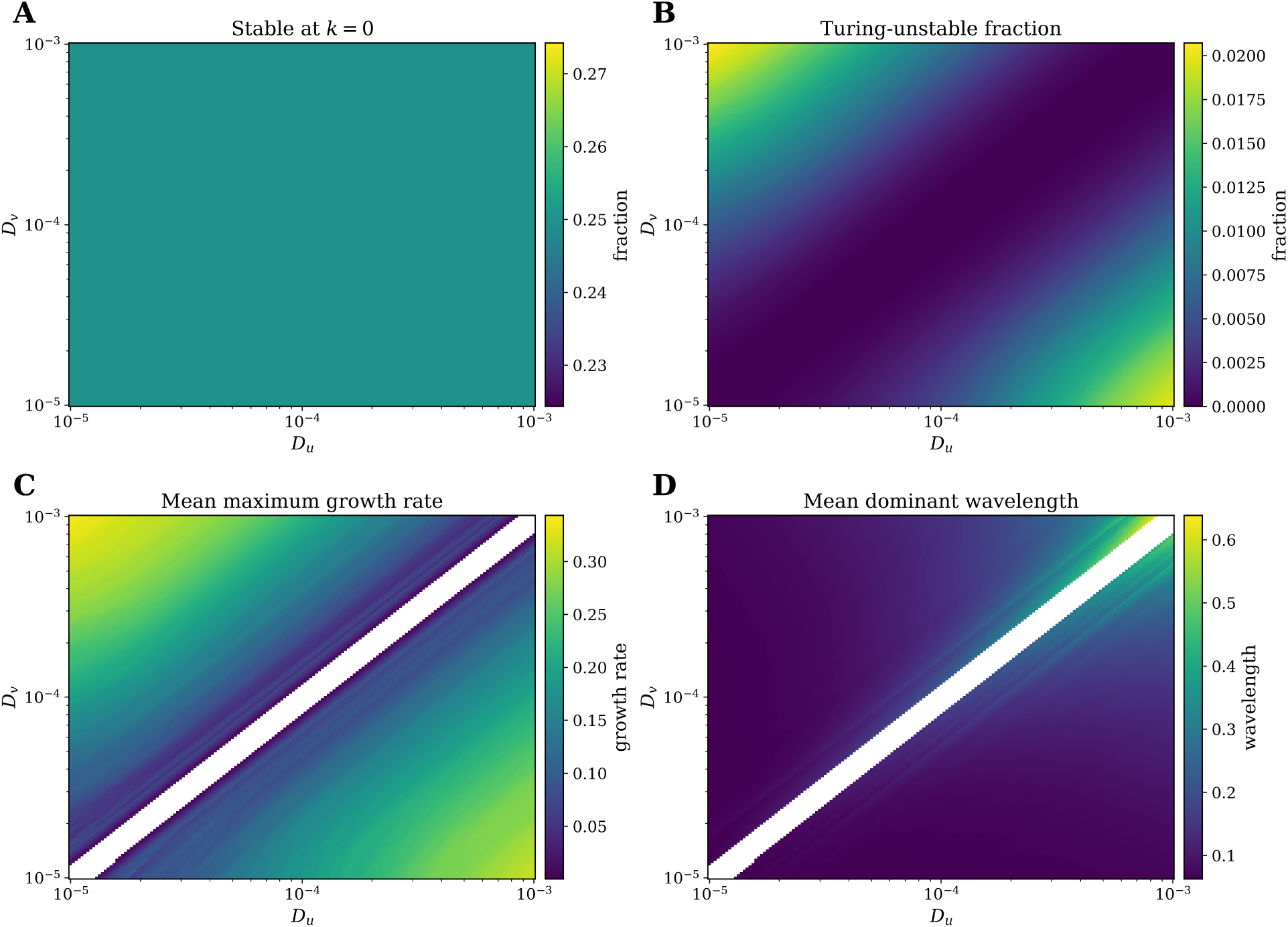
Spectral summary of the linearized reaction–diffusion system. For each diffusion pair (*Du, Dv*), Jacobian entries (*a, b, c, d*) were sampled uniformly from [−2, 2], and the dispersion relation of the linearized two-species system was evaluated over a range of wavenumbers *k*. Panel (A) shows the fraction of sampled Jacobians whose homogeneous steady state is stable at *k* = 0. Panel (B) shows the fraction that are Turing-unstable, that is, stable at *k* = 0 but unstable for some finite *k >* 0. Panel (C) shows the mean maximum growth rate ⟨max*_k>_*_0_ max*_i_* ℜ*λ_i_*(*k*)⟩ across the Turing-unstable cases. Panel (D) shows the mean dominant wavelength ⟨2*π/k*max⟩, where *k*max is the wavenumber with maximal growth rate. White regions in panels (C) and (D) indicate diffusion pairs for which no sampled Jacobian satisfied the Turing instability criterion, so no mean growth rate or dominant wavelength is defined.

The stability analysis provides a spectral description of how perturbations around a homogeneous steady state behave. It identifies regions of parameter space where diffusion-driven instabilities occur, the growth rates of the unstable modes, and the characteristic wavelengths selected by the linear dynamics. In this sense it explains where spatial structure can first emerge, but only at the level of local behavior near the steady state.

At the same time, the information obtained from the dispersion relation is necessarily limited. It does not reveal the form of the realized patterns once nonlinear effects become significant, nor does it quantify how diverse, structured, or irregular those patterns become. Two parameter sets with similar linear instability properties can still produce very different spatial organizations, and instability itself does not guarantee that persistent patterns will survive at long times. In some cases, spatial structure may be only transient before the system returns to a homogeneous state.

These limitations motivate the use of complementary descriptors that act directly on the patterns produced by the full system. In the next section, the Diversity of Number of States (DNOS) and the Diversity of Pattern Complexity (DPC) are introduced as the primary operational measures used in this study to quantify realized patterns across models and parameter regimes. The Uniformity Measure Entropy (UME) is used more narrowly as an auxiliary normalized descriptor of uniformity that helps interpret or refine these summaries where appropriate. Together, these measures extend the linear stability picture by connecting instability structure to the diversity and organization of the patterns that actually form.

### 2.2 Operational DNOS–DPC measurement framework

This section defines the operational measurement framework used throughout the manuscript. Rather than assigning patterns to geometric taxonomies (e.g. “spots” versus “stripes”), we quantify each realized field using two primary coordinates: the Diversity of Number of States (DNOS) and the Diversity of Pattern Complexity (DPC). DNOS measures effective state-space occupancy, or equivalently how broadly a pattern explores its available intensity or alphabet space. DPC measures per-pattern structural complexity through lossless compression of a fixed digital representation. Both coordinates are operational rather than universal: their numerical values depend on the stated normalization, discretization, serialization, and compression protocol. This dependence is intentional and controlled. The framework is therefore used as a reproducible comparative coordinate system, not as a fundamental complexity law. Figure 2 provides an illustrative overview of how DNOS and DPC are constructed; formal definitions and implementation details are given in Section 4.3 (see in particular Sections 4.4–4.6).

**Figure 2:**
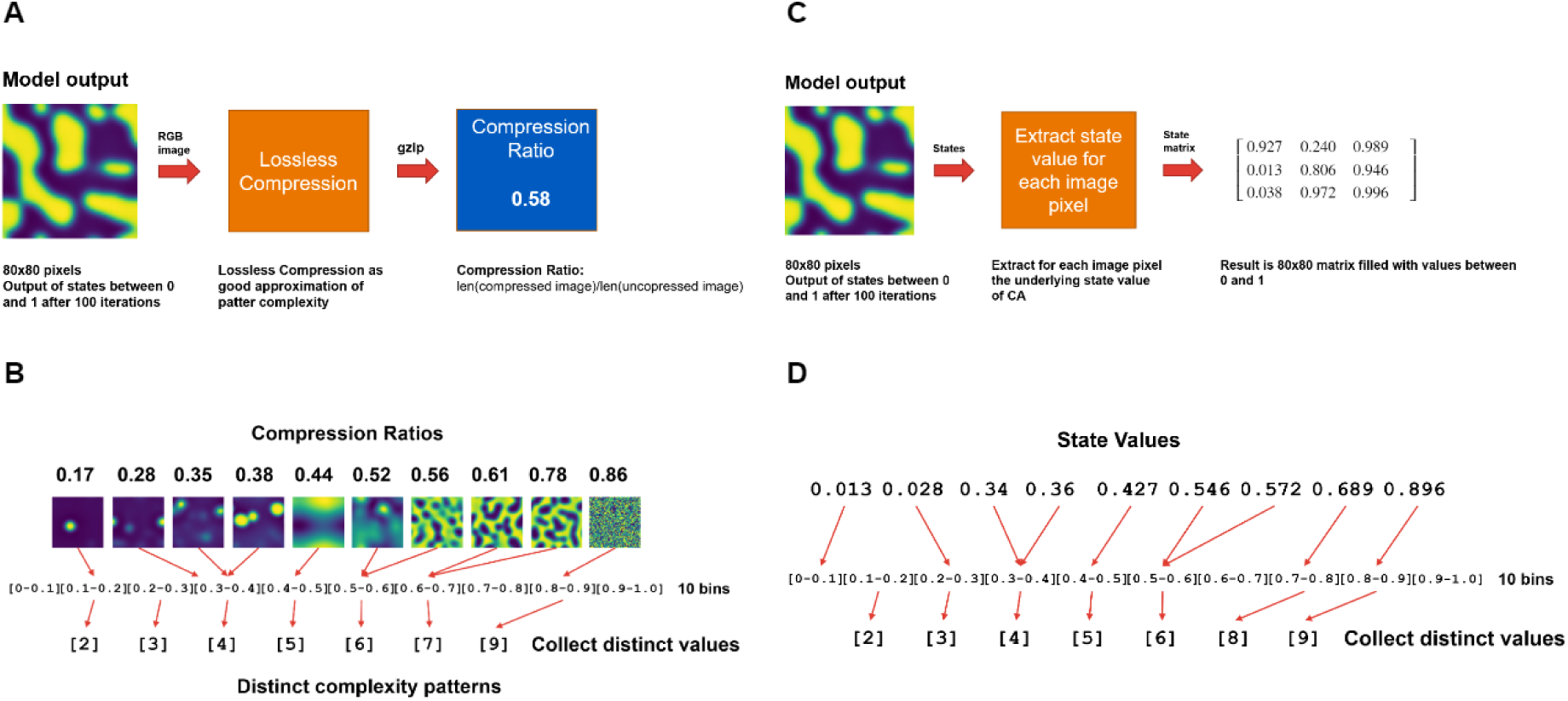
Schematic construction of the complexity measures. **A,B:** Illustration of the Diversity of Pattern Complexity (DPC). Each pattern is rendered as an image and compressed using a lossless codec (gzip), yielding a per-pattern compression ratio DPC(*I*) = |gzip(*I*)|/|*I*|. For the schematic we group these ratios into coarse bins to show how complexity scores are distributed across a collection of patterns. **C,D:** Illustration of the Diversity of Number of States (DNOS). A normalized model output on an 80 × 80 grid with values in [0, 1] is discretized into a small number of equal-width bins (here *B* = 10 for visual clarity), and each pixel is mapped to its bin index; the set of occupied bins provides an intuitive notion of how many intensity levels are realized. In the Methods (Section 4.4) this occupied-bin definition is evaluated at a much finer resolution *B*^∗^ and compared with an entropy-based DNOS diagnostic to verify consistency.

The construction of DNOS begins by normalizing the model output to the unit interval, *X*(**x**) ∈ [0, 1], on a finite pixel grid (typically 80 × 80 in the schematic example). In the toy illustration of Fig. 2C,D we discretize [0, 1] into a small number of equal-width bins (here *B* = 10), and map each pixel value to its bin index. For example, a state value of 0.013 falls into the [0.0, 0.1) bin, while a value of 0.34 falls into the [0.3, 0.4) bin. The set of distinct bin indices occupied by the pattern then encodes, at a coarse level, how many intensity intervals are actually used. In the Methods (Section 4.4) we retain this *occupied-bin* definition as our operational DNOS, evaluated at a high fixed bin count *B*^∗^, and show that it aligns with a more classical entropy-based definition when *B* is sufficiently large. In this sense, DNOS should be interpreted as a measure of realized alphabet breadth or state-space occupancy at a controlled observational resolution.

The second primary measure, DPC, quantifies structural complexity via lossless compressibility. Starting from the same pixel grid, we quantize the normalized field to a fixed integer representation and compress the resulting byte string (gzip in the main text), obtaining for each *individual* pattern *I* a compression ratio

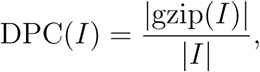

as detailed in Section 4.5. Low DPC values correspond to highly compressible, regular, or nearly homogeneous patterns, whereas high DPC values indicate less compressible and therefore structurally richer configurations under the same representation. Panels 2A,B use a coarse binning of these per-pattern DPC scores to illustrate how compression ratios are distributed across a collection of patterns, but throughout the paper DPC is treated as a *per-pattern* scalar measure and used directly, without bin-counting, in the regime-identification pipeline.

As a secondary supporting descriptor, we also introduce the Uniformity Measure of Entropy (UME) as a summary of how evenly a pattern occupies its available intensity (or complexity) space (Section 4.6). Given a binned distribution over *B* equally spaced intensity bins, with empirical probabilities *p_i_*, UME is defined as the entropy normalized by its maximum possible value,

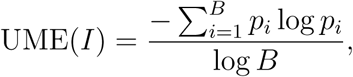

taking the value 1 for perfectly uniform histograms and 0 when all mass is concentrated in a single bin. We apply UME primarily to intensity histograms, providing a measure of how uniformly intensity space is explored and thereby supporting the interpretation of DNOS, and, in auxiliary analyses, to distributions of DPC scores. UME therefore serves as an auxiliary bridge to more conventional entropy-based summaries rather than as a co-equal headline complexity measure.

Taken together, DNOS and DPC define the core comparative space used throughout this study: DNOS captures how many effective states are realized, and DPC captures how structurally complex the resulting spatial configurations are. UME is used only as a supporting interpretive quantity when needed to clarify how evenly those states or complexity levels are distributed.

Because DNOS and DPC are protocol-dependent, we explicitly tested the sensitivities that are most likely to affect interpretation: histogram resolution for DNOS, digital encoding and compressor choice for DPC, modest observational noise, and clustering method for regime identification. These checks are summarized in Figs. S2–S4 and Sec. S5 of the supplementary material. In brief, DNOS is reported at a fixed high-resolution bin count after comparison with entropy-based and bin-free diversity references; DPC rankings are conserved across lossless compression settings and remain stable under modest image perturbations; and the main regime partitions are stable to clustering method and bin-count variation. These robustness analyses do not make the measures protocol-free. Instead, they justify using the explicitly stated DNOS–DPC protocol as a stable comparative coordinate system for the model ensembles analyzed below.

### 2.3 Exploring the complexity space of the linearized reaction–diffusion system

Having characterized the spectral behavior of the linearized system, we now examine how the same parameter space is organized when patterns are described through their realized spatial structure. The complexity measures introduced in the preceding section, based on the diversity of occupied intensity states and the compressibility of the final spatial configuration, allow us to evaluate, for each parameter set, the degree of variation and structural richness present in the patterns that emerge after the system evolves beyond the initial linear stage. This approach addresses a complementary question to the stability analysis. Whereas the dispersion relation identifies parameter combinations that permit the growth of spatial modes, the complexity-based analysis quantifies the patterns that are actually produced once the system has evolved for a finite time in the fully discretized numerical model. The two descriptions therefore capture different aspects of the same system: the linear analysis predicts the onset and preferred wavelength of spatial structure, while the complexity measures summarize the organization of the resulting patterns across the entire parameter ensemble.

Applied to the linearized Turing model, this dual perspective reveals that the presence of instability is not by itself predictive of the eventual diversity or detail of the patterns. Instead, the complexity measures identify distinct regions in the diffusion plane, with each region associated with characteristic levels of state diversity and structural organization. This organization is not fully visible from the spectral data alone but becomes clear when DPC and DNOS are examined together.

In this section we numerically solve the PDE system using the finite-difference method (Section 4.10) and revisit the linearized reaction–diffusion system by exploring a comprehensive range of diffusion and reaction parameters to investigate their impact on pattern formation in two-dimensional concentration fields. The parameters and their respective ranges are shown in Table 1.

**Table 1:** Summary of the parameters and their ranges explored in the linearized Turing simulations.

| Parameter | Description | Range | Increment |
| --- | --- | --- | --- |
| $D_u$ (log scale) | Diffusion rate of $u$ | $10^{-3}$ to $10^{-5}$ | 0.2 (logarithmic) |
| $D_v$ (log scale) | Diffusion rate of $v$ | $10^{-3}$ to $10^{-5}$ | 0.2 (logarithmic) |
| $a$ | Reaction rate for $u$ | $-2$ to $2$ | 0.2 |
| $b$ | Interaction of $v$ with $u$ | $-2$ to $2$ | 0.2 |
| $c$ | Reaction rate for $u$ | $-2$ to $2$ | 0.2 |
| $d$ | Reaction rate for $v$ | $-2$ to $2$ | 0.2 |
| $dx$ | Spatial resolution | Computed as $1/\text{SIZE}$ | |
| $dt$ | Time step | 0.02 | |

In our analysis, we focus on the simulation outputs after 10^3^ time steps for all combinations of diffusion coefficients (*D_u_, D_v_*) and reaction parameters *a*, *b*, *c*, and *d*. For each parameter combination, we compute DPC from the gzip compression ratio of the *u*-field snapshot (Methods, Section 4.5). To summarize how complexity varies across the (*D_u_, D_v_*) grid, we discretize the resulting DPC values to four decimal places (equivalently, into uniform bins at that resolution) and count the number of distinct values observed across all (*a, b, c, d*) combinations for that (*D_u_, D_v_*) pair.

For DNOS, we similarly discretize the *u*-field concentration values to four decimal places for every reaction-parameter combination, count all distinct values, and average the resulting occupied-bin counts across (*a, b, c, d*). This procedure is equivalent to evaluating the DNOS_occ_ measure defined in the Methods (Section 4.4) at a fixed high intensity resolution. Thus each point in the (*D_u_, D_v_*) plane carries a pair of coarse-grained complexity descriptors derived from per-pattern DPC and DNOS values.

We next examine the organization of these two descriptors by plotting all normalized DPC and DNOS values for all (*D_u_, D_v_*) pairs in the feature plane (Figs. 3C–F). All clustering in this study is performed using *k-means*, consistent with the implementation used for every model and figure in the paper. (As discussed in Methods, Section 4.8, Gaussian Mixture Models and hierarchical clustering yield essentially identical partitions.)

**Figure 3:**
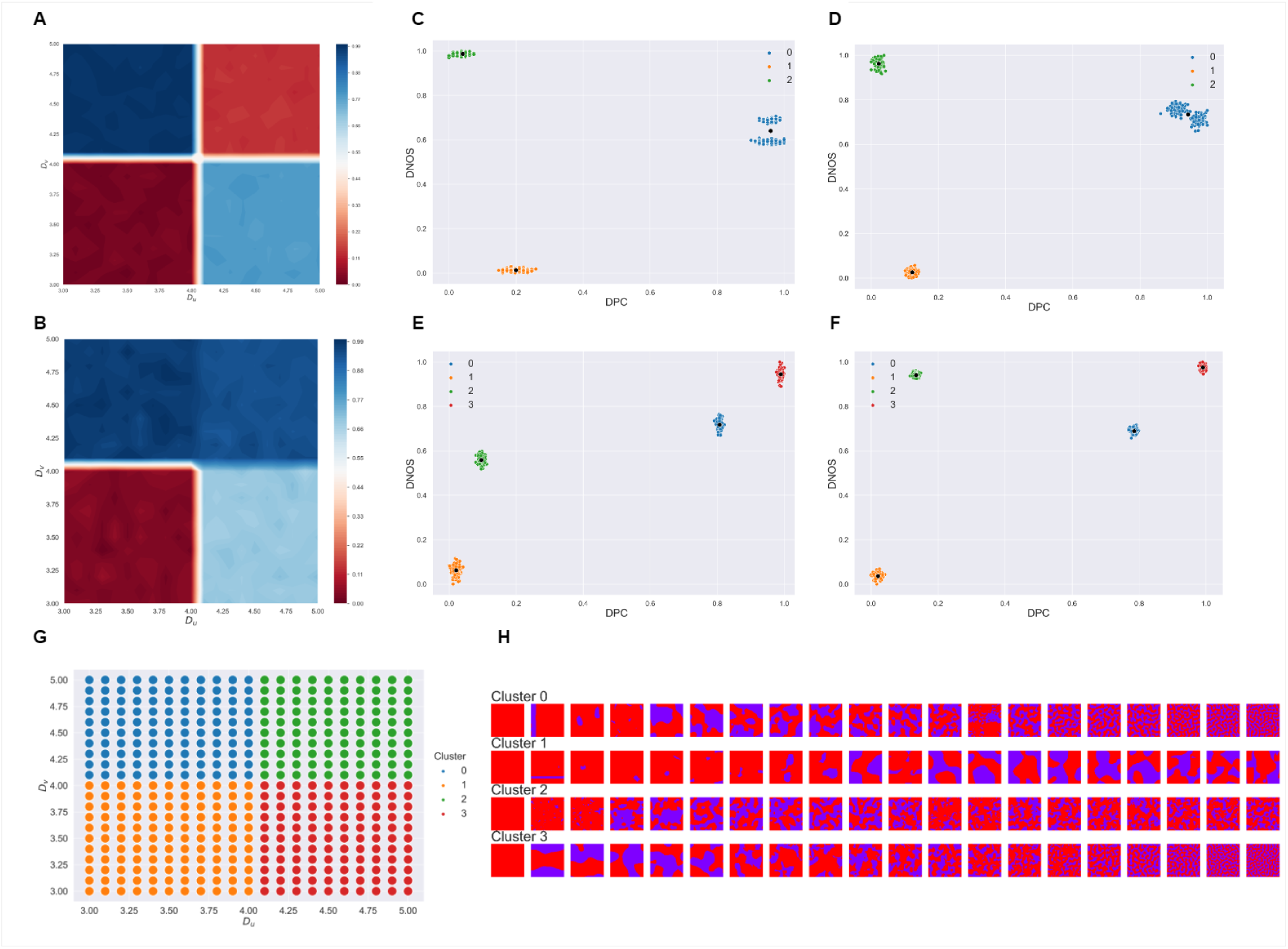
Exploring the complexity space of the linearized reaction–diffusion system. **A,B:** Contour plots of normalized DPC-and DNOS-based summaries over the (*Du, Dv*) plane (Methods, Sections 4.5–4.4). **C–F:** Scatter plots of normalized DPC and DNOS for all (*Du, Dv*) pairs under two effective resolutions. All clustering shown uses *k-means*; alternative clustering algorithms give equivalent partitions (Methods, Section 4.8). Panels **E,F** incorporate UME (Methods, Section 4.6) as an additional feature, producing sharper separation of regimes. **G:** Mapping cluster assignments from **F** onto the (*Du, Dv*) plane. **H:** Representative patterns per cluster, sorted by DPC, illustrating the characteristic variability within each regime. All complexity measures are computed from final-time patterns after transient decay, and the (*Du, Dv*) plane is sampled on a refined uniform grid to avoid artifacts from coarse parameter sampling.

To illustrate sensitivity to resolution we show an effective binning into 10^3^ bins (Fig. 3C) and 10^4^ bins (Fig. 3D). In both cases, four clear clusters emerge. A lower-left cluster (label 1) exhibits low DPC and low DNOS; an upper-left cluster (label 2) has low DPC but high DNOS; and two upper-right clusters (labels 0 and 3) correspond to high DPC and high DNOS. At the coarser resolution the points appear smeared along the DPC axis, whereas the finer resolution produces more compact clusters.

To refine separation, particularly in the high-DPC/high-DNOS corner where cluster 0 appears split, we apply the Uniformity Measure of Entropy (UME; Methods, Section 4.6) before clustering. UME distinguishes patterns with similar bin occupancy but different degrees of uniformity. Including UME yields four sharply separated clusters for both resolutions (Figs. 3E,F), with the high-complexity corner cleanly separating into clusters 0 and 3.

Figures 3A,B show contour plots of the normalized DPC-based and DNOS-based summaries over the (*D_u_, D_v_*) plane. Both reveal quadrant-like segmentation, particularly pronounced in the DPC landscape. The lower-left and upper-right quadrants correspond to low complexity, while the upper-left quadrant exhibits the highest complexity.

We then map the k-means cluster assignments back onto the (*D_u_, D_v_*) grid (Fig. 3G), showing that the four clusters align cleanly with the diffusion quadrants identified in the contour plots. Clusters 1 and 2 predominantly occur along the diagonal, while clusters 0 and 3 occupy off-diagonal regions.

Finally, to visualize the qualitative patterns underlying each cluster, we extract representative patterns (Fig. 3H). For each (*D_u_, D_v_*) assigned to a given cluster, we collect its patterns, scale and discretize them, identify mode patterns in DPC bins, and sample 20 images evenly across the resulting complexity range. Cluster 1 exhibits the lowest diversity (nearly homogeneous states); Cluster 2 shows greater diversity despite low DPC; and Clusters 0 and 3 display rich, multi-scale structures, with Cluster 3 showing the greatest variation across scales.

These results establish the central point of the framework: the onset of instability does not by itself determine the realized organization of biological patterns, whereas DNOS and DPC provide a direct comparative description of that realized structure.

### 2.4 The Gray–Scott model

The Gray–Scott model involves two reacting and diffusing chemical species that produce rich stationary and dynamic spatial patterns. Although the Gray–Scott kinetics were originally introduced as a simplified autocatalytic scheme rather than a literal biochemical pathway [28], the model has become a prototypical example of nonlinear reaction–diffusion dynamics exhibiting spots, stripes, labyrinths, and self-replicating structures (e.g. [29–31]).

Extensive mathematical and computational analyses have clarified the conditions under which these patterns form and persist, highlighting multistability, hysteresis, and complex localized states such as pulses and self-replicating spots [32–34]. As a result, the Gray–Scott model is widely used as a benchmark system for studying pattern formation in chemical, biological, and ecological contexts.

We write the Gray–Scott dynamics in the standard form

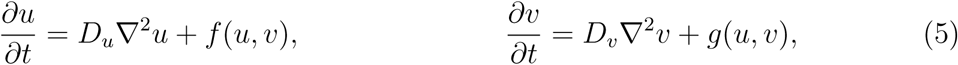

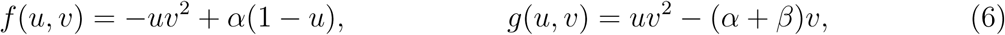

where *u* and *v* are the concentrations of the two species, *D_u_*and *D_v_*are their diffusion coefficients, ∇^2^ is the Laplacian, and *α*, *β* control feed and removal rates. This representation fits naturally into the general Turing framework introduced earlier.

We collect outputs after 10^2^ time steps for all combinations of diffusion coefficients (*D_u_, D_v_*) and reaction parameters (*α, β*) (Table 2).

**Table 2:** Summary of the parameters and their ranges explored in the Gray–Scott simulations.

| Parameter | Description | Range | Increment |
| --- | --- | --- | --- |
| $D_u$ | Diffusion rate of $u$ | 0.1 to 0.2 | 0.005 |
| $D_v$ | Diffusion rate of $v$ | 0.01 to 0.1 | 0.0045 |
| $\alpha$ | Feed rate | 0 to 0.01 | 0.0005 |
| $\beta$ | Removal rate | 0 to 0.01 | 0.0005 |
| $dx$ | Spatial resolution | 1 | |
| $dt$ | Time step | 1 | |

For each parameter combination, we compute DPC from the gzip compression ratio of the *u*-field snapshot and DNOS from the occupied-bin count of the normalized intensity histogram.

Figures 4A,F show contour plots of normalized DPC and DNOS over the (*D_u_, D_v_*) plane. To benchmark DPC against established quantitative descriptors of spatial complexity, we also compute three auxiliary measures: Fourier spectral energy (ESP), wavelet energy ratio (WLE), and the resistance-distance histogram (RDH) (Sec. S6 of the supplementary material). Their corresponding landscapes are shown in Figs. 4B–D, and pairwise Pearson correlation coefficients (PCC) and mean squared errors (MSE) are summarized in Fig. 4E. DPC shows especially strong agreement with RDH and WLE, indicating that it tracks established notions of structural complexity while remaining simple and mechanism-agnostic under the fixed representation and compression protocol used here.

**Figure 4:**
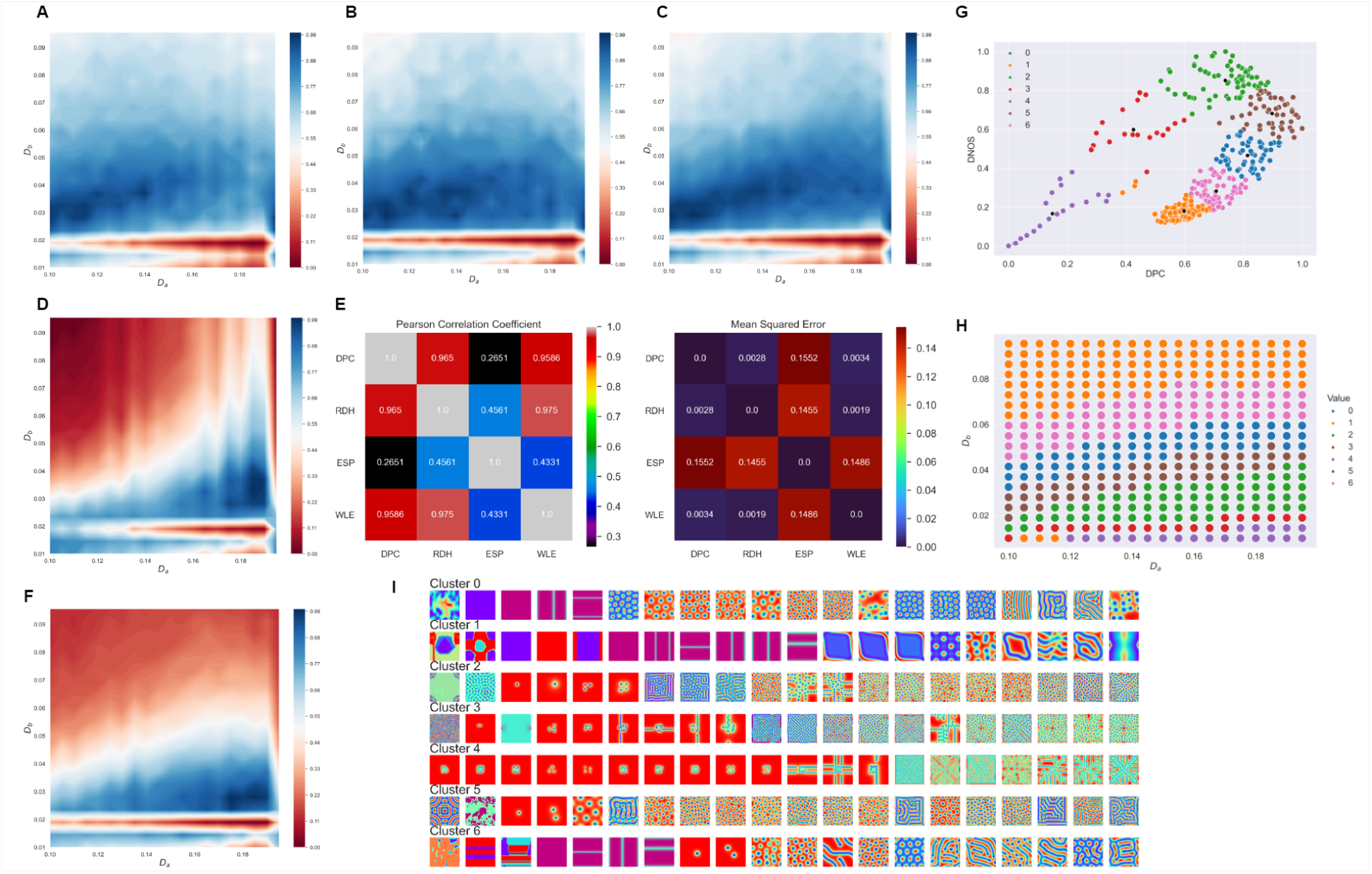
The Gray–Scott model. **A,F:** Contour plots of normalized DPC and DNOS, respectively, over the *Du* × *Dv* plane. **B–D:** Contour plots of the auxiliary benchmarking measures wavelet energy ratio (WLE), resistance-distance histogram (RDH), and Fourier spectral energy (ESP), respectively, over the same parameter plane (Sec. S6 of the supplementary material). **E:**Pairwise Pearson correlation coefficients and mean squared errors (MSE) between the normalized DPC, RDH, ESP, and WLE arrays, shown as correlation and MSE heatmaps. **G:** Scatter plot of normalized DPC versus DNOS for all (*Du, Dv*) pairs, with *k*-means cluster assignments indicated by color. **H:** Mapping of the *k*-means clusters from panel **G** back onto the *Du* × *Dv* plane. **I:** For each cluster, 20 representative patterns selected around the cluster centroid, sorted by DPC and arranged in a grid, illustrating the diversity of spatial phenotypes within each regime.

We next plot all normalized DPC and DNOS values for all (*D_u_, D_v_*) pairs in the feature plane (Fig. 4G). In contrast to the clean quadrant structure observed there (Fig. 3C–F), the Gray–Scott feature space shows more intricate regions, with clusters forming bands and patches spread across the (*D_u_, D_v_*) plane. This indicates that some diffusion parameter combinations support markedly higher pattern complexity and diversity than others. Mapping these cluster assignments back onto the (*D_u_, D_v_*) grid (Fig. 4H) shows that the parameter space is partitioned into distinct bands associated with more or less complex regimes.

Representative samples around the cluster centroids, sorted by DPC (Fig. 4I), visualize the range of spatial phenotypes associated with each computational regime in the Gray–Scott model. Taken together, these results show that DNOS and DPC remain informative beyond the linear benchmark, recovering structured nonlinear regimes in a canonical reaction–diffusion system.

### 2.5 The FitzHugh–Nagumo model

Having shown in the Gray–Scott system that DNOS and DPC recover structured nonlinear regimes beyond the linear benchmark, we next ask whether the same framework remains informative in a mechanistically different reaction–diffusion model with excitable dynamics. The FitzHugh–Nagumo (FHN) model is a reduced form of the Hodgkin–Huxley equations [35], capturing neuronal excitability through a two-variable system introduced by FitzHugh [36] and Nagumo [37]. Its simplicity and canonical cubic nonlinearity make it a standard test case in reaction–diffusion theory, where it supports traveling waves, oscillations, and Turing-like spatial structure [38, 39].

Following [38], we use a reaction–diffusion formulation

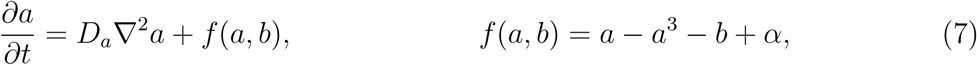

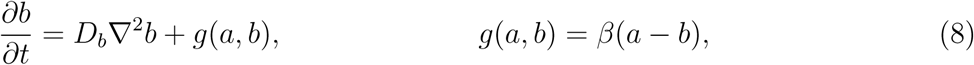

where *a* acts as an activator and *b* as an inhibitor. The cubic term *a* − *a*^3^ captures excitable nonlinearity, while −*b* and *β*(*a* − *b*) provide inhibition and coupling. Diffusion coefficients *D_a_, D_b_* and parameters *α, β* control transitions between uniform, oscillatory, and patterned states.

We simulated the system across the full parameter grid listed in Table 3, collecting final snapshots after 10^2^ time steps.

**Table 3:** Parameter ranges explored in the FitzHugh–Nagumo simulations.

| Parameter | Description | Range | Increment |
| --- | --- | --- | --- |
| $D_u$ | Diffusion rate of $a$ | 1 to 50 | 2.5 |
| $D_v$ | Diffusion rate of $b$ | 1 to 100 | 5 |
| $\alpha$ | Activator bias | 0 to 0.1 | 0.005 |
| $\beta$ | Coupling rate | 0 to 0.1 | 0.005 |
| $dx$ | Spatial resolution | 1 | |
| $dt$ | Time step | 0.001 | |

We constructed contour landscapes of normalized DPC and DNOS over the (*D_u_, D_v_*) plane (Fig. 5A,F). In parallel, we computed three auxiliary descriptors, the resistance-distance histogram (RDH), Fourier spectral energy (ESP), and wavelet energy ratio (WLE), as defined in Sec. S6 of the supplementary material, to benchmark DPC against established pattern–complexity measures (Fig. 5B–D).

**Figure 5:**
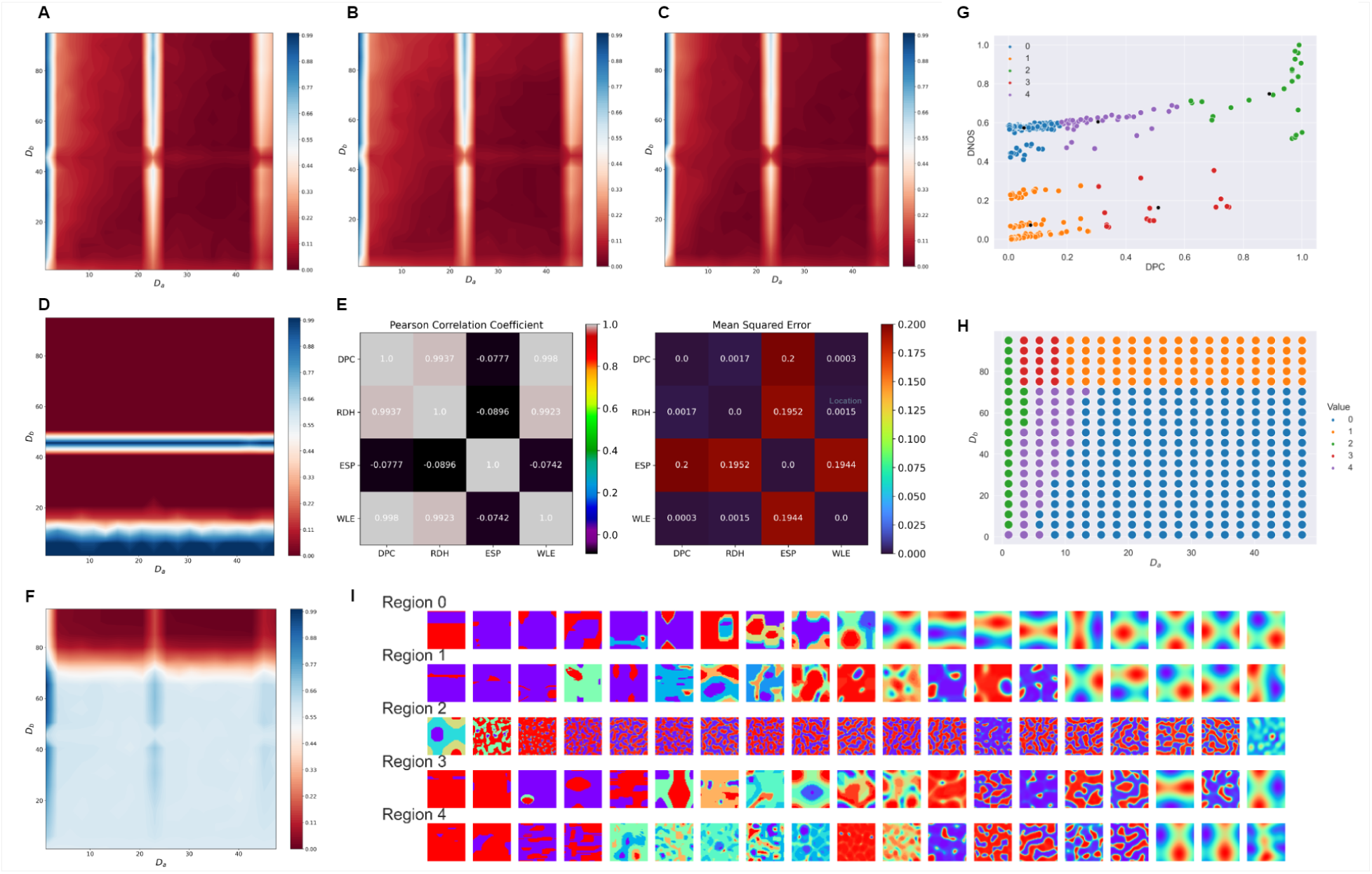
The FitzHugh–Nagumo model. **A,F:** Contour plots of normalized DPC and DNOS, respectively, over the *Du* × *Dv* plane. **B–D:** Contour plots of the auxiliary benchmarking measures wavelet energy ratio (WLE), resistance-distance histogram (RDH), and Fourier spectral energy (ESP), respectively, over the same parameter plane (Sec. S6 of the supplementary material). **E:** Pairwise Pearson correlation coefficients and mean squared errors (MSE) between the normalized DPC, RDH, ESP, and WLE arrays, shown as correlation and MSE heatmaps. **G:** Scatter plot of normalized DPC versus DNOS for all (*Du, Dv*) pairs, with *k*-means cluster assignments indicated by color. **H:** Mapping of the *k*-means clusters from panel G back onto the *Du* × *Dv* plane. **I:** For each cluster, 20 representative patterns selected around the cluster centroid, sorted by DPC and arranged in a grid, illustrating the diversity of spatial phenotypes within each regime.

Pairwise Pearson correlations and mean squared errors (MSE) between the normalized measures are shown in Fig. 5E. DPC exhibits extremely high correlations with RDH (*ρ* = 0.994) and WLE (*ρ* = 0.998), together with very low MSE values (DPC–RDH: 0.00166, DPC–WLE: 0.00029). These near–perfect concordances indicate that DPC captures structural and topological detail almost identically to multiscale (WLE) and graph-based (RDH) descriptors. By contrast, DPC shows weak and slightly negative correlation with ESP (*ρ* ≈ −0.20), reflecting that spectral energy emphasizes a different dimension of complexity, as already observed for the Gray–Scott model (Section 2.4).

Clustering in the DNOS–DPC plane identifies structured regimes within the FitzHugh–Nagumo parameter space (Fig. 5G,H). Representative patterns sampled around the cluster centroids (Fig. 5I) show that these regimes correspond to distinct realized spatial phenotypes, spanning simpler states and markedly more complex multiscale configurations. Taken together, the Gray–Scott and FitzHugh–Nagumo results show that the DNOS–DPC framework is not tied to one specific nonlinear mechanism, but instead provides a common comparative description across distinct classes of reaction–diffusion dynamics.

### 2.6 Spatiotemporal patterning enabled by gene regulatory networks

Gene regulatory networks (GRNs) are central to spatiotemporal pattern formation in biological systems, governing cell–fate specification, boundary formation, and multicellular organization [40, 41]. Among the canonical GRN motifs is the *toggle switch*, comprising two mutually repressing genes, which generates bistability and enables cells to commit robustly to distinct transcriptional states [42–44]. A prominent example is the GATA1–PU.1 switch in hematopoiesis, which stabilizes lineage commitment by mutual antagonism of master regulators.

When embedded in a spatially extended medium with 2D diffusion, the toggle switch can generate stable spatial partitions, pinned domain walls, and heterogeneous steady states [45–47]. Adding positive autoregulation introduces additional attractors (including tristability), increasing both dynamical richness and pattern diversity [48, 49]. Such extensions are relevant for tissue development, lineage branching, and pathological dysregulation such as malignant dedifferentiation [50].

We next ask whether the same comparative framework remains informative when spatial organization is shaped primarily by regulatory architecture rather than by canonical activator–inhibitor kinetics. To address this, we analyze two related systems introduced in Roy et al. [51]: (i) a toggle switch with 2D diffusion only, and (ii) a toggle switch with 2D diffusion plus self-activation. Their comparison tests whether DNOS and DPC can detect how changes in regulatory motif architecture alter realized spatial organization.

#### 2.6.1 Toggle switch with 2D diffusion

The toggle-switch dynamics follow

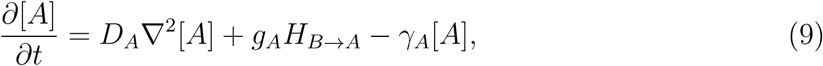

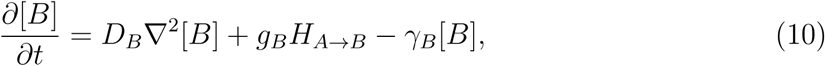

where *H_X_*_→*Y*_ denotes Hill-type inhibition,

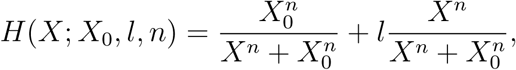

with threshold *X*_0_, fold-change *l*, and Hill coefficient *n*. Diffusion coefficients, degradation rates, and production rates are sampled across the ranges listed below, following [51].

We simulated the system on a 50 × 50 grid (5 mm × 5 mm), with *u* and *v* initialized to 100. Diffusion was set to *D* = 0.001, and integration was performed for 500 time steps with *dt* = 0.1. Parameters were varied over: Hill coefficients *n_A_*_→_*_B_, n_B_*_→_*_A_* ∈ [1, 6] (step 0.25), thresholds *A*_0*B*_, *B*_0*A*_ ∈ [1, 10] (step 1), and ten random draws of (*γ, g*) for each pair of Hill exponents.

Contour plots of normalized DPC and DNOS across this parameter grid are shown in Fig. 6A,B. DPC exhibits diagonal bands of high and low complexity, indicating alternating parameter regimes in which diffusion and bistability promote or suppress spatial patterning.

**Figure 6:**
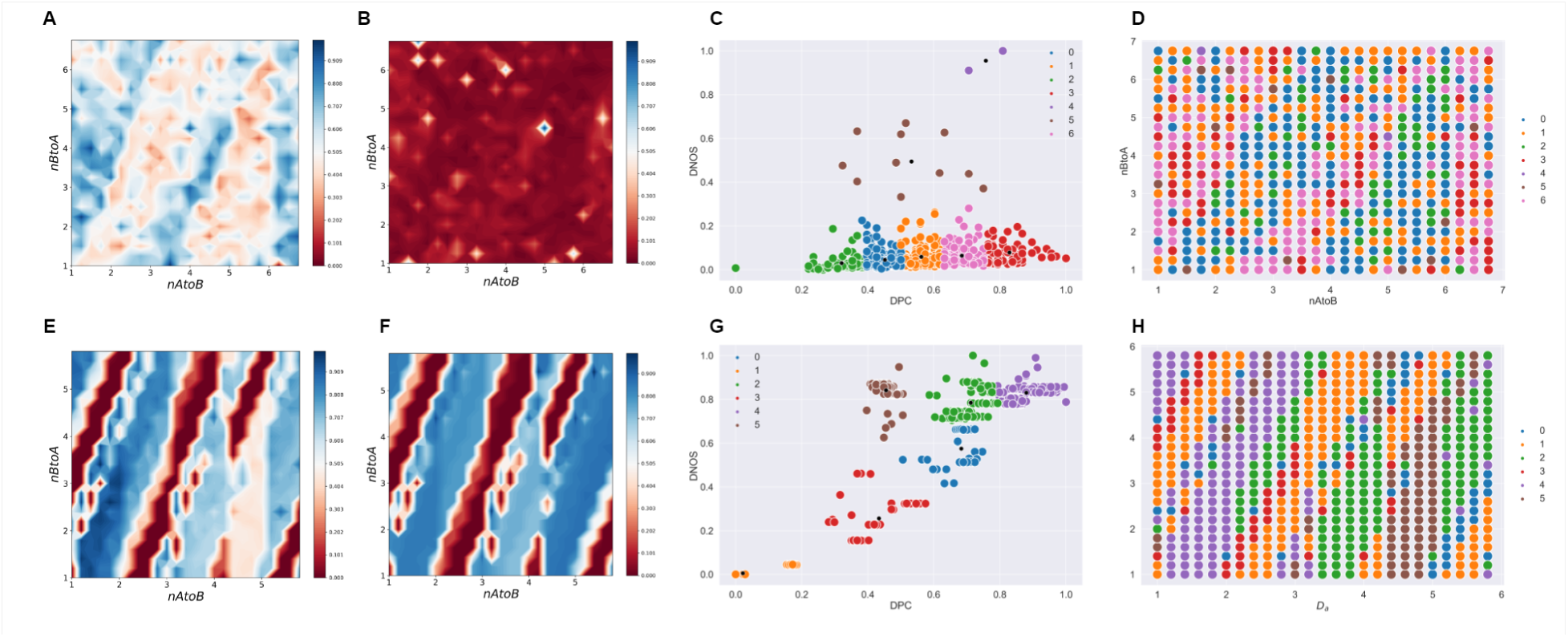
Toggle switch models. **A,B:** Normalized DPC and DNOS landscapes for the toggle switch with 2D diffusion. **C:** DPC–DNOS scatter with k-means clusters. **D:** Cluster assignments mapped back to the parameter grid. **E,F:** Normalized DPC and DNOS for the toggle switch with 2D diffusion and self-activation. **G:** DPC–DNOS scatter with k-means clusters for the self-activating system. **H:** Clusters mapped to parameter space.

By contrast, DNOS remains uniformly low across nearly the entire space, suggesting that although patterns differ in compressibility (DPC), they occupy only a narrow subset of intensity space and thus exhibit limited state diversity.

Clustering in the DNOS–DPC plane shows that most solutions remain in a low-DNOS regime while spanning a broad range of DPC values (Fig. 6C,D). Thus, the diffusion-only toggle switch generates substantial variation in structural complexity while remaining constrained in realized state diversity.

#### 2.6.2 Toggle switch with 2D diffusion and self-activation

Adding positive autoregulation modifies the PDEs to

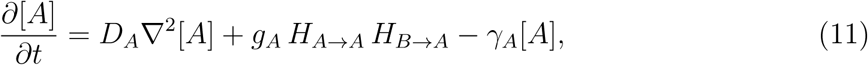

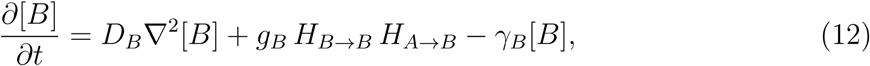

with *H_A_*_→*A*_ and *H_B_*_→*B*_ representing self-activation (Hill-type positive feedback), and with the same Hill-function form as above.

Simulations were performed on a 20 × 20 grid (2 mm × 2 mm) for 5000 time steps, with diffusion coefficient *D* = 10^−5^. Parameters were sampled across: *n_A_*_→_*_A_, n_A_*_→_*_B_* ∈ [1, 6] (step 0.2), thresholds *A*_0*A*_, *A*_0*B*_ ∈ [1, 6] (step 0.25), fold changes *λ_A_*_→_*_A_* = 3.0, *λ_A_*_→_*_B_* = 0.1, and random draws of (*γ, g*) following [51].

Contour plots of normalized DPC and DNOS (Fig. 6E,F) reveal markedly richer structure than in the non-self-activating case. DPC again forms diagonal bands, but the transitions between high- and low-complexity regions are much more sharply defined. Unlike the 2D-diffusion-only model, DNOS now also separates into distinct regions of high and low diversity, indicating that self-activation enables the system to explore a significantly larger effective state space.

In the self-activating system, DNOS and DPC increase together much more clearly than in the diffusion-only toggle switch (Fig. 6G,H), indicating that positive feedback expands both realized state diversity and structural complexity. Viewed together, the two toggle-switch models show that DNOS and DPC are sensitive not only to parameter variation within a model, but also to qualitative changes in regulatory architecture.

Unlike the FitzHugh–Nagumo system, which is excitable and supports traveling pulses, the toggle-switch systems considered here produce stationary domain walls and pinned interfaces governed by bistability or tristability. These mechanistic differences lead to distinct structures in the (DPC, DNOS, UME) feature space, despite superficial similarities in the visual appearance of some patterns.

### 2.7 Cross-model comparison in the DNOS–DPC–UME space

Before turning to the developmental application, we summarize how the different model classes analyzed above occupy a common complexity space using UME as an auxiliary descriptor of uniformity (Sec. S4 of the supplementary material). DNOS and DPC define the primary comparative axes of the framework, while UME indicates how evenly patterns explore their realized state or complexity ranges. For each system, we aggregate UME computed on (i) per-pattern compression ratios, UME(CR), and (ii) normalized state values across space, UME(*S*), yielding one summary point per model in the UME(CR), UME(*S*) plane.

Figure 7 shows the resulting positions for the five systems studied in this work: the linearized Turing model (LINTP; Section 2.3), the Gray–Scott model (GSM; Section 2.4), the FitzHugh–Nagumo model (FHN; Section 2.5), the toggle switch with 2D diffusion (TGLS; Section 2.6.1), and the toggle switch with 2D diffusion and self-activation (TGLSWA; Section 2.6.2).

**Figure 7:**
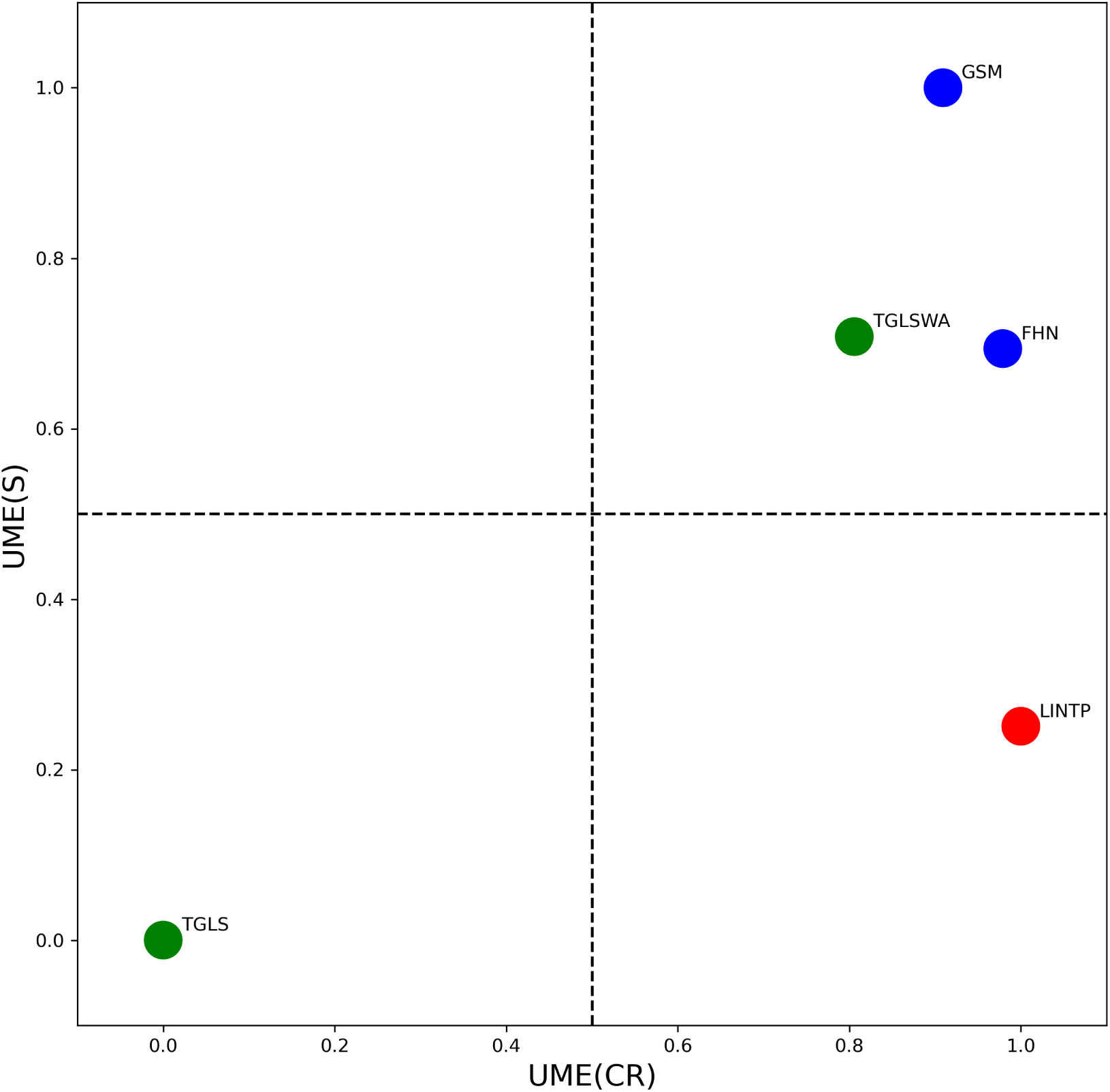
Cross-model comparison in the UME summary space. Each point shows the Uniformity Measure of Entropy applied to compression ratios (UME(CR), horizontal axis) and to state values (UME(S), vertical axis) for one of the five models: linearized Turing (LINTP), Gray–Scott (GSM), FitzHugh–Nagumo (FHN), toggle switch with 2D diffusion (TGLS), and toggle switch with 2D diffusion and self-activation (TGLSWA). Different mechanistic classes occupy distinct regions of this common complexity space, allowing their realized spatial organization to be compared within a single operational framework.

Across all systems, the joint DNOS–DPC–UME representation is consistent with the main comparative picture developed above: different mechanistic models occupy distinct regions of feature space, yet remain describable within a common operational framework. In particular, Gray–Scott, FitzHugh–Nagumo, and the self-activating toggle switch occupy comparatively high-uniformity regions, whereas the linearized benchmark and the diffusion-only toggle switch are more restricted. This cross-model summary therefore provides the clearest overall comparison of how reaction–diffusion and gene-regulatory systems realize spatial structure across mechanistic classes.

### 2.8 Notch, Delta, and EGF signaling in *Drosophila* neurogenesis

Finally, we test whether the DNOS–DPC framework remains biologically informative in a concrete developmental system, using the Notch–Delta–EGF model of *Drosophila* neurogenesis. Classical Notch–Delta lateral inhibition produces “salt-and-pepper” patterns during binary cell-fate decisions, but in the medulla a distinct traveling “proneural wave” propagates across the neuroepithelium. This wave is driven by proneural transcription factors such as Lethal of scute and is modulated by an EGF-mediated reaction–diffusion process that suppresses the usual lateral-inhibition pattern and instead establishes a coherent front of differentiation [52, 53]. In this system Notch does not simply impose a static spatial alternation pattern. Rather, EGF diffusion regulates the velocity and spatial organization of Notch activation, highlighting a mode of GRN-based pattern formation that depends on coupled reaction–diffusion and lateral-inhibition dynamics.

We analyze the reaction–diffusion GRN model introduced by Sato et al. [52], focusing on the four interacting fields: the EGF ligand *E*, the Notch receptor *N*, the Delta ligand *D*, and the proneural factor *A*. Our aim is not to reinterpret the underlying biological mechanism established by that work. Instead, we use the model as a mechanistically grounded test case for a complementary measurement question: once the coupled EGF–Notch–Delta–proneural dynamics have unfolded, how is realized spatial organization distributed across molecular fields, the full signaling state, and developmental routes? Their dynamics are governed by

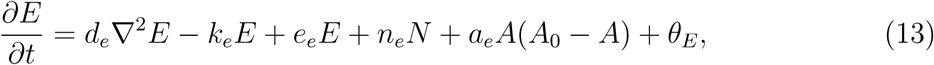

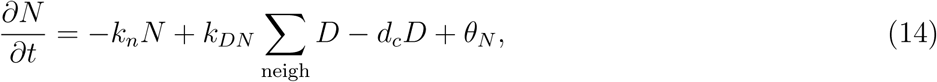

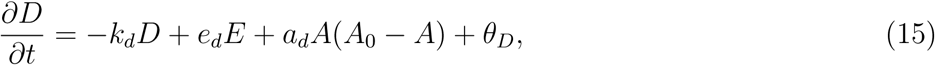

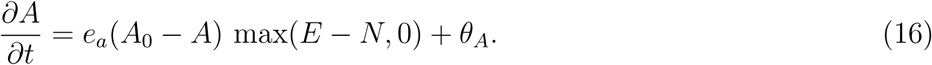

Here ∇^2^ denotes the Laplacian; *d_e_* is the only diffusing species, as in the experimental system; *k_DN_* denotes neighbour-mediated Delta-to-Notch coupling and is distinct from the numerical timestep Δ*t*; and the remaining interactions encode mutual reinforcement and inhibition between *E*, *N*, *D*, and *A*. Stochastic terms *θ_E,N,D,A_* represent molecular noise.

To simulate mutant clones, individual regulatory interactions are set to zero following Sato et al. [52]. In EGF-deficient clones, the self- and *A*-mediated activation of *E* (*e_e_* and *a_e_*) are suppressed. In Notch mutants, the terms required for Notch signaling are removed. In Delta mutants, Delta-production and Delta-mediated signaling terms are suppressed. This gate-based intervention isolates the contribution of each pathway to pattern formation while preserving the same dynamical framework for all genotypes.

We solve the PDE system using the finite-difference scheme described in Section 4.10. We deliberately retain the static final-snapshot endpoint analysis as the first level of the Drosophila result, because it provides continuity with the original DNOS–DPC comparison and with the biological question of whether mutant clones reach different final patterning regimes. We then use a focused expanded analysis to ask what lies behind that endpoint shift: which fields carry the loss or persistence of organization, whether the full multichannel signaling state separates genotypes more clearly than any single field, and whether mutants follow different routes through complexity space before reaching their endpoints.

The endpoint comparison uses the three parameter sets in Table 4. The expanded field-resolved, multichannel, and time-resolved trajectory analysis uses parameter set C, because this set fixes the core biochemical scales while sampling the interaction strengths most directly associated with proneural-wave propagation. Simulations were run for wild type, EGF, Delta, and Notch conditions on a 20 × 20 grid for 2000 iterations, with 1000 random parameter samples per genotype. To convert the endpoint analysis into a dynamical analysis, the released workflow saves intermediate checkpoints and computes DNOS, DPC, and UME on each field at each checkpoint. For single-field measurements, normalization uses the run-specific range of that channel over all checkpoints; for the multichannel state, the reference range is computed over all channels and checkpoints for the same run. Endpoint statistics are reported for the final checkpoint, while trajectory statistics are computed from the full ordered sequence of checkpointed values. The expanded analysis should therefore be read as a controlled extension of the original final-snapshot comparison, not as a replacement for it: Fig. 8 asks where final states lie, whereas Figs. 9–12 ask which fields carry the separation and how each genotype reaches its final region.

**Figure 8:**
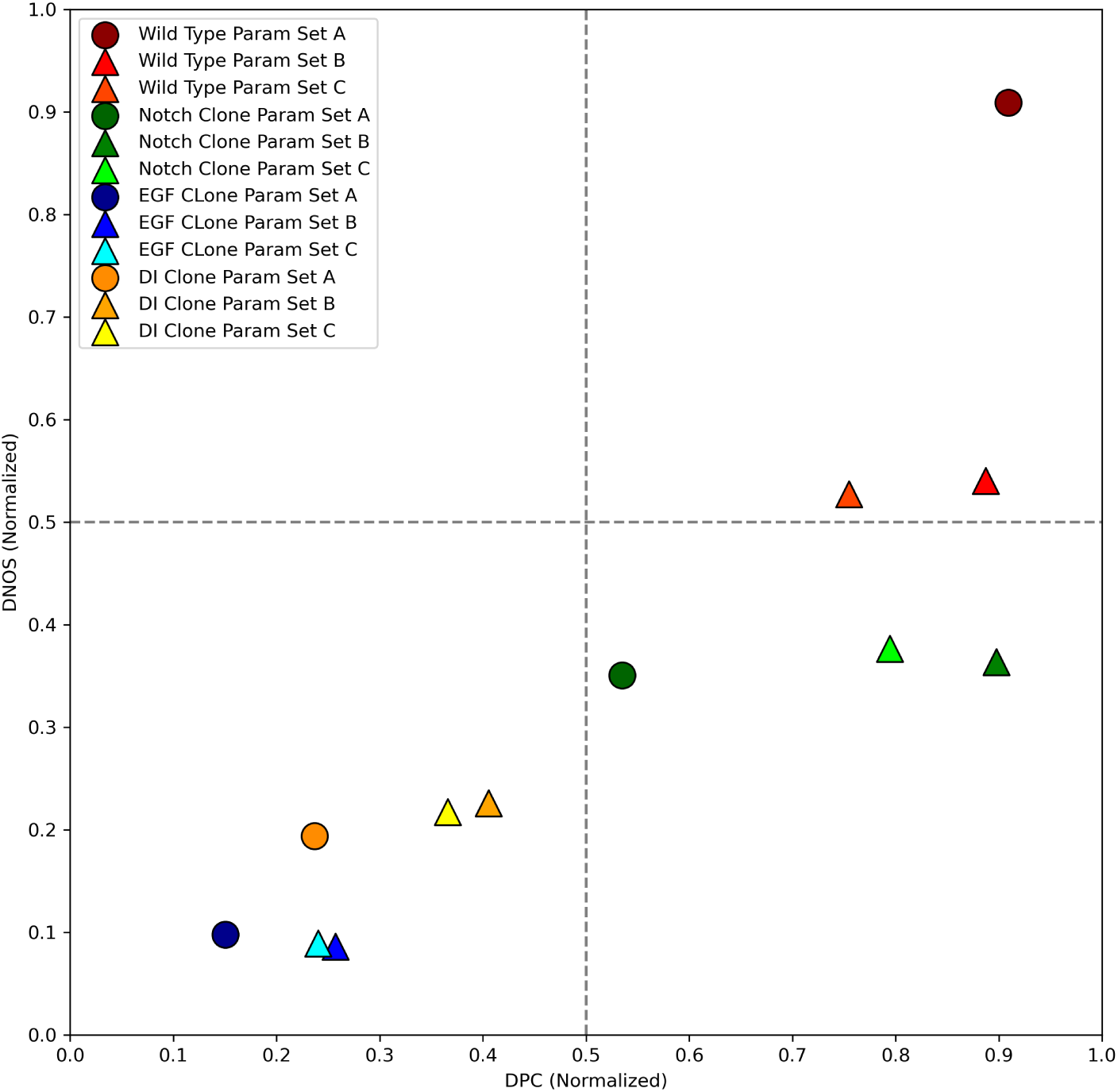
Static endpoint DNOS–DPC landscape in Notch–Delta–EGF signaling. Normalized pattern-compression complexity (DPC) plotted against the normalized diversity of number of states (DNOS) for final wild-type and EGF, Notch, and Delta mutant simulations. Points are color-coded by clone type and marker shapes indicate the three parameter sets A–C. Dashed lines at 0.5 on both axes divide the space into four coarse regions. Wild-type simulations occupy the high-complexity/high-diversity region, whereas mutant endpoint clouds shift toward lower DPC and/or DNOS values under this final-snapshot protocol. This figure establishes endpoint genotype separation, but it does not by itself identify whether the separation reflects upstream pathway collapse, downstream proneural-output persistence, or altered developmental routes. Those distinctions are addressed in Figs. 9–12.

**Figure 9:**
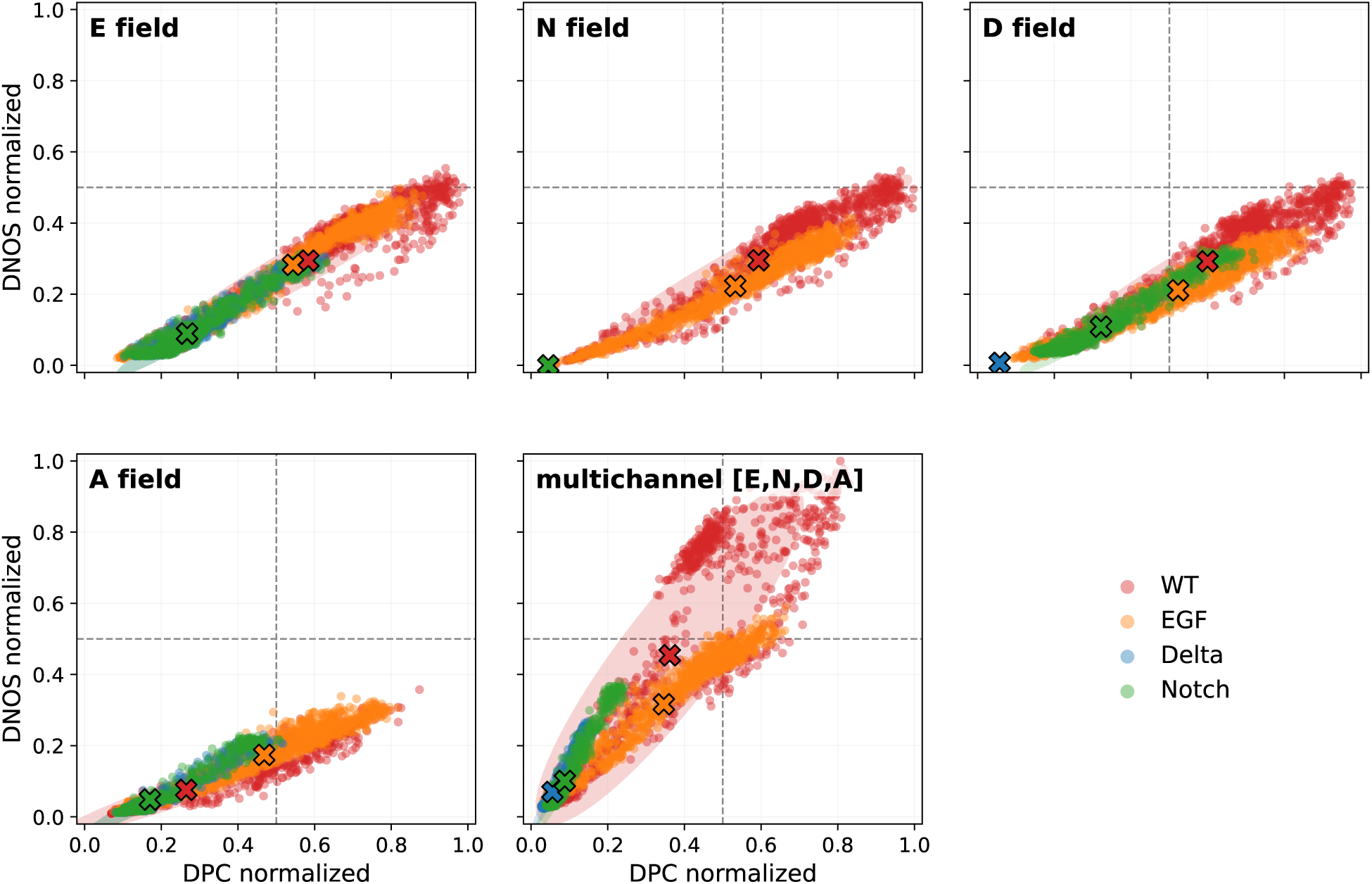
Field-resolved final DNOS–DPC landscapes in the Notch–Delta–EGF model. Normalized DPC and DNOS are shown for the final checkpoint of each wild-type, EGF, Delta, and Notch simulation. Separate panels show the *E*, *N*, *D*, and *A* fields, together with the multichannel signaling state [*E, N, D, A*]. Points show individual parameter samples, cross markers show genotype means, and shaded ellipses summarize genotype-level covariance. The multichannel panel provides the clearest integrated endpoint separation, with wild type occupying the highest joint DNOS–DPC region and EGF mutants showing broad reduction of the coupled signaling state. The *N*-field panel identifies pathway-level collapse under Delta/Notch disruption, whereas the *A*-field shows that downstream proneural-state organization can persist even when upstream signaling channels are degraded. Thus an organized *A*-field endpoint is not interpreted as proof of normal development unless the upstream fields and trajectory are also considered.

**Figure 10:**
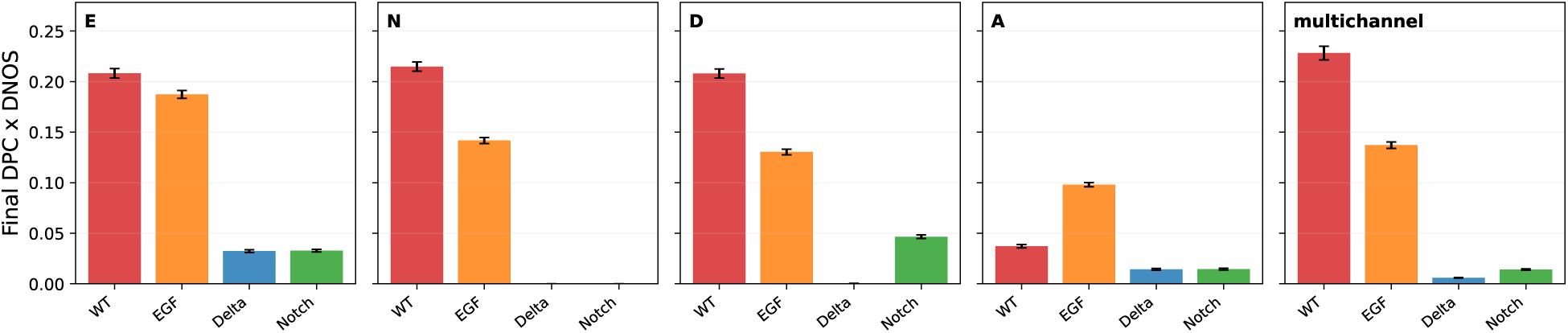
Endpoint organization score across fields. For each genotype and field, the figure shows the mean final product DPC × DNOS with standard error across parameter samples. This product is used as a compact display statistic, not as a replacement for the two primary axes. Wild type remains high in the *E*, *N*, *D*, and multichannel readouts, whereas mutant effects are field-specific. The *A*-field is not interpreted as a monotonic measure of correct wave progression; elevated or persistent *A*-field organization can reflect downstream proneural heterogeneity even when upstream signaling is disrupted. Delta and Notch therefore show selective pathway deficits rather than a uniform loss of complexity.

**Figure 11:**
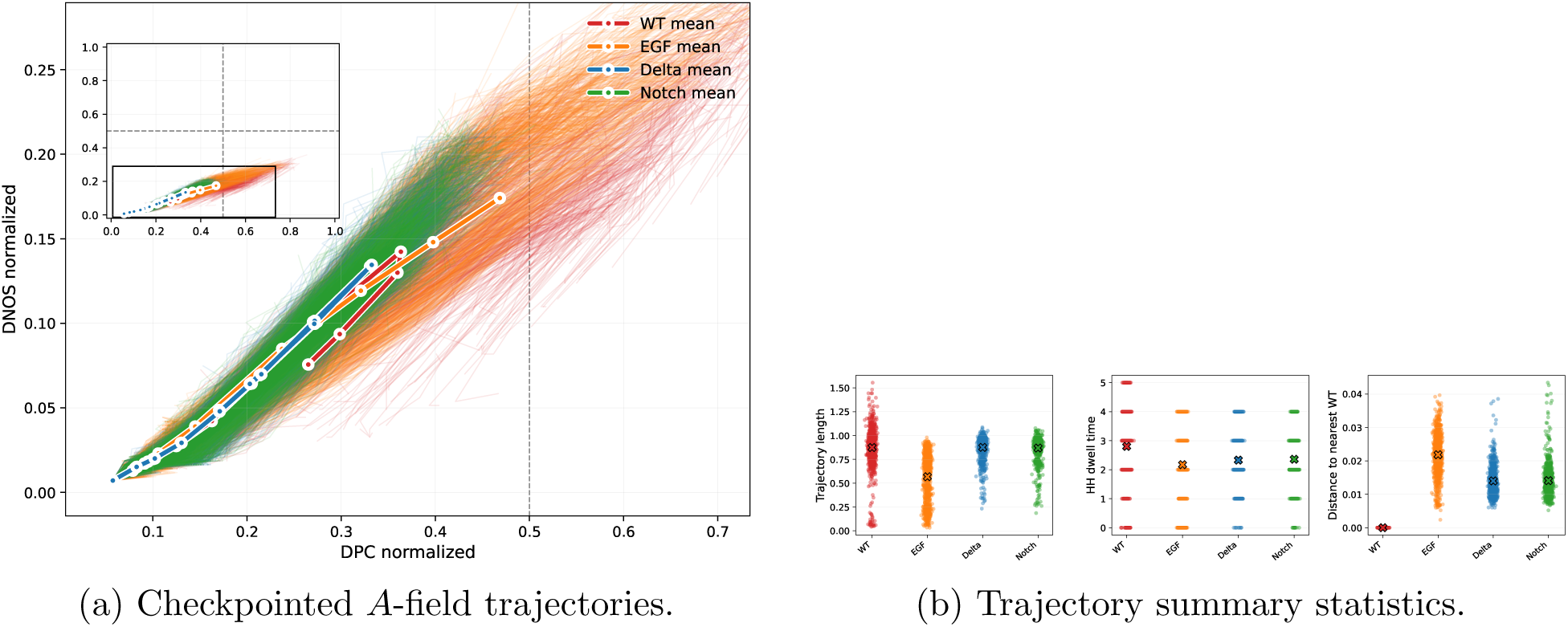
Time-resolved DNOS–DPC analysis of proneural *A*-field dynamics. (A) Each simulation defines a trajectory through the DNOS–DPC plane as the proneural field develops over checkpointed time. Thin curves show individual runs and thick curves show genotype means. (B) Trajectory length, high-organization dwell time, and distance to the nearest wild-type trajectory quantify genotype-specific developmental routes. EGF mutants show the strongest reduction in trajectory length and high-organization dwell in this ensemble, consistent with an entry/progression defect into organized proneural dynamics. Delta and Notch retain longer *A*-field trajectories despite field-specific disruption of upstream signaling, indicating downstream persistence rather than globally normal patterning or full rescue of wild-type development.

**Figure 12:**
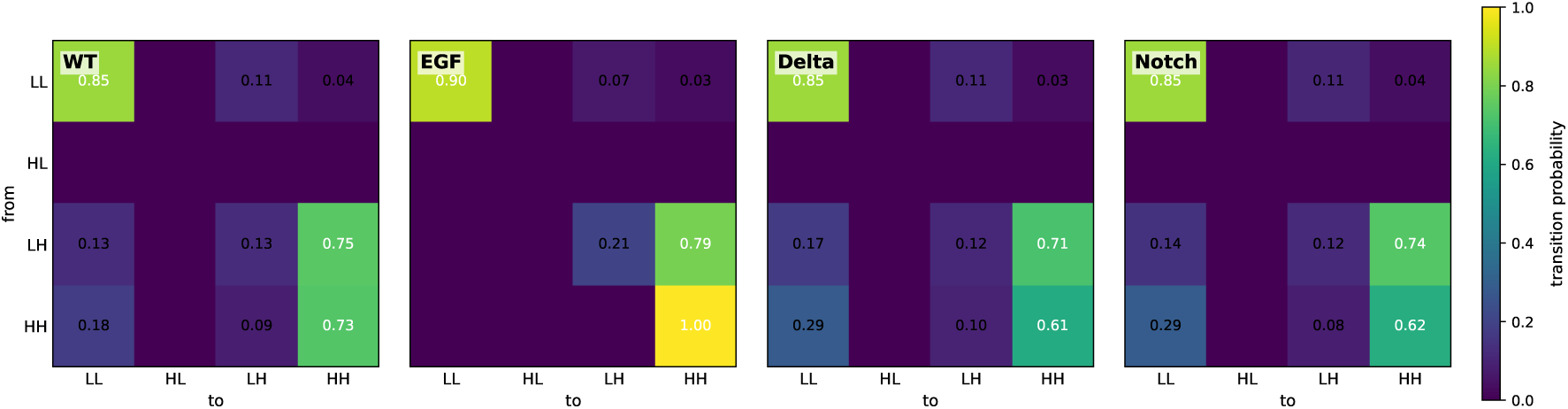
Coarse-grained *A*-field regime transitions. Checkpointed trajectories were coarse-grained into LL, HL, LH, and HH regimes using pooled upper-quartile thresholds in the normalized DNOS–DPC plane; the first letter denotes DNOS status and the second denotes DPC status. Entries show transition probabilities between consecutive checkpoint regimes. EGF mutants show an entry/progression bias, with stronger LL retention and reduced escape from the low-organization regime. Delta and Notch mutants can reach organized *A*-field states but show weaker HH self-persistence and greater leakage back to LL than wild type. These transition matrices provide a compact state-machine representation of developmental progression through complexity space.

**Table 4:** Combined Parameter Settings for Proneural Wave Experiments.

| Parameter | Set A | Set B | Set C |
| --- | --- | --- | --- |
| $d_e$ | [0.0, 1.0] | [0.0, 1.0] | Fixed at 1.0 |
| $e_d$ | [0.0, 1.0] | [0.0, 1.0] | Fixed at 1.0 |
| $A_0$ | [0.0, 1.0] | [0.0, 1.0] | Fixed at 1.0 |
| $n_e$ | [0.0, 1.0] | [0.0, 1.0] | [0.0, 2.0] |
| $a_d$ | [0.0, 1.0] | [0.0, 1.0] | Fixed at 1.0 |
| $k_n$ | [0.0, 1.0] | [0.0, 1.0] | Fixed at 1.0 |
| $k_e$ | [0.0, 1.0] | [0.0, 1.0] | Fixed at 1.0 |
| $k_d$ | [0.0, 1.0] | [0.0, 1.0] | Fixed at 1.0 |
| $e_e$ | [0.0, 1.0] | [0.0, 1.0] | [0.0, 0.4] |
| $a_e$ | [0.0, 1.0] | [0.0, 1.0] | [0.0, 2.0] |
| $e_a$ | [0.0, 10.0] | [0.0, 10.0] | Fixed at 10.0 |
| $k_{DN}$ | Fixed at 0.01 | [0.0, 1.0] | [0.0, 0.5] |
| $d_c$ | Fixed at 0.25 | [0.0, 1.0] | [0.0, 0.5] |

For each final-snapshot simulation in the original endpoint analysis we computed DNOS and DPC using the same binning, normalization, and compression procedure as in the preceding model classes. DNOS and DPC values were min–max normalized within the Notch–Delta–EGF endpoint ensemble to allow direct comparison across wild-type and mutant systems. Figure 8 shows the resulting static endpoint DNOS–DPC landscape across parameter sets A–C and clone types. In this endpoint representation, wild-type points occupy the high-DPC/high-DNOS region, indicating both high structural organization and broad state occupancy, whereas the mutant endpoint clouds shift toward lower DNOS and/or lower DPC. This is the appropriate conclusion from Fig. 8: final realized patterns differ systematically between wild type and the perturbations. It is not, by itself, a claim that lower DNOS–DPC is a universal measure of biological abnormality, nor that all mutants fail by the same mechanism.

Importantly, the endpoint plot alone cannot determine whether each mutant changes realized organization by the same mechanism. It also cannot distinguish loss of upstream pathway organization from persistence, broadening, or reorganization of the downstream proneural output. We therefore treat the static landscape in Fig. 8 as the baseline genotype-level summary and then ask, in the expanded analysis below, which biological fields and which temporal routes account for this separation.

We next decomposed this endpoint result by field and by developmental route. For each checkpointed simulation we computed DNOS and DPC separately for each biological field *E*, *N*, *D*, and *A*, and also for the multichannel signaling state [*E, N, D, A*]. The single-field analysis asks which molecular or regulatory component carries structural complexity.

The multichannel analysis asks how complex the full tissue signaling state is when all four variables are considered jointly. All measures were computed with the same bin count and compression procedure used elsewhere in the paper, then normalized over the complete Notch–Delta–EGF analysis so that field-specific and multichannel panels can be compared on a common DNOS–DPC scale.

Figure 9 shows the resulting field-resolved endpoint landscapes for the expanded analysis. These panels refine, rather than replace, the static endpoint conclusion in Fig. 8. The key biological point is that the four fields do not carry the same information. The *A*-field endpoint captures the downstream proneural-output state and therefore reports whether a proneural pattern has been realized, but it does not by itself identify which signaling module produced that output or whether the route was wild-type-like. Delta and Notch perturbations can overlap partially with wild type in the *A*-field; this is not a contradiction of the endpoint separation, but evidence that downstream proneural organization can persist or reorganize despite upstream signaling disruption. By contrast, the *N*-field gives a more pathway-proximal readout: Delta and Notch perturbations collapse the Notch-signaling field toward very low DNOS and DPC, whereas wild type maintains a structured *N*-field. The *E*- and *D*-field panels provide complementary readouts of reaction–diffusion propagation and ligand-mediated lateral signaling. The clearest integrated separation appears in the multichannel [*E, N, D, A*] representation, where wild-type simulations occupy the highest joint DNOS–DPC region, EGF mutants show a broad reduction in the coupled signaling state, and Delta and Notch mutants occupy intermediate, field-specific regimes. Thus the expanded analysis changes the biological interpretation of the endpoint landscape: mutant phenotypes are not simply weaker versions of wild type, but distinct redistributions and losses of organization across different channels in the coupled EGF–Notch–Delta–proneural system.

To quantify these endpoint effects, we summarized the genotype means of final DPC, final DNOS, and their product DPC × DNOS across fields. The product is not introduced as a new primary measure; it is used only as a compact visual summary of joint structural complexity and state diversity. Figure 10 shows that wild type is highest in the *E*, *N*, *D*, and multichannel readouts, whereas EGF, Delta, and Notch perturbations reduce different parts of the signaling state. The *A*-field should be interpreted separately: endpoint *A*-field organization is not a monotonic proxy for correct proneural-wave progression, because a stalled, broadened, or heterogeneous proneural output can still occupy a relatively large DNOS–DPC region. Thus the field summary is most informative when read together with the time-resolved trajectories. Biologically, this is consistent with EGF loss weakening the reaction–diffusion component that supports wave propagation through the coupled signaling state, Delta removing lateral signaling input, and Notch disruption collapsing the Notch-signaling channel while allowing partial downstream proneural organization in some parameter regimes.

We next asked whether genotype differences are visible only at the endpoint or whether they also alter the developmental route through complexity space. For each run and field we defined a checkpointed trajectory

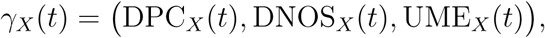

where *X* ∈ {*E, N, D, A,* [*E, N, D, A*]}. Trajectory length was computed in this three-dimensional coordinate system, while endpoint positions and regime labels were evaluated in the normalized DNOS–DPC plane. We then computed trajectory length, the first entry into a high-organization regime, the dwell time in that regime, and the time-aligned distance to the nearest wild-type trajectory. Regimes were defined by coarse-graining the normalized DNOS–DPC plane using pooled upper-quartile thresholds: LL denotes low DNOS and low DPC, HL high DNOS and low DPC, LH low DNOS and high DPC, and HH high DNOS and high DPC.

Figure 11 shows the *A*-field trajectories and their summary statistics. All genotypes move from low-DNOS/low-DPC initial states toward more organized regions, consistent with progressive proneural-wave development. However, the route, not only the endpoint, is genotype dependent. In this ensemble, EGF mutants have the shortest trajectories, reduced dwell time in high-organization regimes, and the largest distance from wild-type trajectories. This pattern is consistent with an EGF-dependent entry or progression defect: without the reaction–diffusion component, simulations are less able to move the proneural field through the typical sequence of increasing structural complexity and state occupancy. Delta and Notch mutants retain longer *A*-field trajectories and can enter organized *A*-field regimes, but the field-resolved endpoint analysis shows that this downstream persistence occurs alongside selective collapse of the upstream Notch-signaling channel. DNOS–DPC trajectories therefore help separate a wave-propagation defect from a lateral-signaling defect that still permits partial proneural-output organization in the downstream field. This is why the trajectory summaries should not be read as a simple complexity ranking: the same downstream DNOS–DPC region can be approached by different signaling routes.

The coarse-grained trajectory transitions provide a final view of these developmental routes. The transition matrices should be interpreted conditionally and route-wise, not as absolute biological cell-state labels. For EGF mutants, the dominant pattern is not an absolute inability to occupy HH once HH has been reached; rather, it is a bias in access to higher-organization regimes. EGF trajectories show stronger retention in LL and a lower probability of leaving LL directly for higher-organization states, consistent with slowed or stalled proneural-wave progression in this simulated ensemble. In contrast, Delta and Notch trajectories can enter organized *A*-field regimes, but their HH states are less self-persistent than wild type and leak back to LL more often. This suggests a different failure mode: lateral signaling is not required for every instance of downstream *A*-field organization, but in this model it appears to help stabilise and coordinate that organization once it has emerged. The near absence of the HL state in all genotypes further indicates that, for this model and measurement protocol, increases in state diversity and structural complexity are coupled rather than occurring as a high-diversity/low-complexity intermediate. Thus the coarse-grained DNOS–DPC representation converts proneural-wave progression into an interpretable state-machine summary: EGF perturbation biases access to organized states, whereas Notch–Delta interactions appear to support maintenance and coordination of organized proneural-output dynamics.

Viewed as a coarse state machine, the transition analysis adds information that is not available from endpoint plots alone. The rarity of a stable HL regime indicates that broad state occupancy without corresponding spatial organization is not a common intermediate in the *A*-field: in this model, DNOS and DPC tend to increase together as proneural-output organization develops. EGF perturbation primarily changes access to organized states, producing stronger LL persistence and reduced progression toward HH. Delta and Notch perturbations instead show that organized *A*-field states can still be reached, but the field-resolved endpoint analysis reveals that these downstream states occur despite collapse or weakening of the upstream Notch–Delta signaling channel. Thus the same downstream DNOS–DPC region can correspond to biologically different signaling states, and the coarse-grained transition matrix provides a compact mathematical abstraction of route-level developmental differences.

This interpretation is consistent with, but does not constitute independent experimental validation of, the broader proneural-wave literature. Experimental and theoretical work has identified EGFR signaling as a driver of excitable proneural-wave propagation in the optic lobe [54]; subsequent work has shown that Delta-dependent cis-inhibition and intracellular trafficking shape the temporal dynamics of Notch activity behind the wavefront [55]; and recent reviews emphasize that Notch dynamics in Drosophila neural development are strongly context-dependent and often require mathematical models to distinguish pathway function from observed spatial output [53, 56]. Our DNOS–DPC analysis should therefore be read as a compact quantitative summary of realized organization in this established modeling context, not as a replacement for model-specific measurements such as wavefront velocity, signaling peak position, field-specific amplitude, or spatial autocorrelation.

Taken together, Figs. 8–12 should be read as a hierarchy of increasingly resolved analyses. The static endpoint landscape establishes genotype-level separation; the field-resolved and multichannel landscapes show which biological variables carry that separation; the trajectory plots show how the downstream proneural field moves through complexity space; and the transition matrices summarize route-level differences between organizational regimes. This final example shows that the framework is not limited to generic benchmark models. In a mechanistically grounded developmental signaling system, DNOS and DPC provide a complementary measurement layer that separates pathway-level and output-level effects, identifies multichannel organization that is partly hidden in single-field readouts, and converts proneural-wave progression into a quantitative trajectory through realized-organization space. The biological conclusion is therefore not that higher DNOS–DPC is automatically more normal, nor that the method discovers the mechanism of proneural-wave propagation. Rather, the analysis quantifies organizational consequences of the Sato et al. model: EGF perturbation biases entry/progression into organized proneural-wave dynamics, Notch–Delta signaling appears to support maintenance and coordination of organized *A*-field states, and the downstream proneural output can remain partially organized even when upstream signaling complexity is lost.

## 3 Discussion

This work develops a quantitative framework for a general problem in nonlinear pattern formation: how to compare the realized spatial organization produced by different systems once they evolve beyond linear onset. Across reaction–diffusion models and gene-regulatory networks, the core operational coordinates DNOS and DPC summarize complementary aspects of that organization, how broadly a pattern occupies its available state space, and how structurally rich or compressible the resulting configuration is under a fixed digital representation. Together, they provide a compact way to compare systems that may differ strongly in mechanism, dimensionality, or visual phenotype, without relying on model-specific geometric labels such as spots or stripes. The framework is deliberately operational: it provides reproducible coordinates for comparing simulated fields, not an absolute measure of physical or biological complexity. UME plays a supporting role by summarising how evenly the intensity or complexity range is occupied when additional uniformity information is useful. Classical linear stability analysis identifies when a homogeneous steady state becomes unstable, but it does not describe the nonlinear pattern that finally emerges or the degree of complexity in that pattern. Across the examples considered here, the joint organization of DNOS and DPC reveals structure in parameter space that is not visible from the linear theory alone. In the linearized reaction–diffusion system, specific regions of the diffusion parameter plane consistently produce patterns with higher or lower DNOS and DPC values, indicating differences in both state diversity and structural detail.

For the nonlinear systems (Gray–Scott, FitzHugh–Nagumo, and the toggle-switch models), DNOS and DPC separate parameter space into regions associated with distinct behavior. High DPC values occur where the models produce self-replicating structures, labyrinths, oscillatory fronts, or multiple domains. Low DPC values correspond to more homogeneous or weakly organized states. The two measures therefore offer a concise way to compare systems whose patterns differ widely in visual appearance.

We compared DPC with several established descriptors (RDH, ESP, and WLE) to assess how it relates to existing measures of pattern irregularity. DPC shows strong agreement with RDH and WLE, which reflect topological and multiscale detail, and weaker agreement with ESP, which reflects spectral energy. This suggests that DPC captures structural detail that is similar to several classical descriptors while also containing information not fully represented by them.

DNOS plays a complementary role. At high resolution it is sensitive to fine-scale occupancy of intensity space, including subthreshold variation and interface structure. UME can then be used as an auxiliary descriptor to distinguish patterns that occupy the available intensity or complexity range more evenly from those that concentrate values in narrow intervals. In this sense, DNOS and DPC define the main comparative space of the framework, while UME provides additional interpretive resolution where needed.

The gene-regulatory models illustrate how this type of quantitative description connects nonlinear pattern organization to biological interpretation. Parameter regimes that include self-activation in toggle-switch systems produce increased DNOS and DPC, consistent with the additional states expected from positive feedback. In the Notch–Delta–EGF system, the retained static endpoint landscape shows genotype-level separation in final pattern complexity, while the expanded field-resolved and time-resolved analyses show why this separation is biologically non-uniform. Different perturbations affect different channels of the coupled signaling system. EGF loss reduces the organization of the multichannel signaling state and biases entry/progression into organized *A*-field dynamics, consistent with the role of EGF-mediated reaction–diffusion in proneural-wave propagation. Delta and Notch perturbations strongly disrupt the Notch-signaling channel, but the downstream proneural *A*-field can still enter partially organized regimes. The coarse-grained transition matrices add a further mechanistic distinction: conditional on reaching HH, EGF trajectories can remain there, whereas their main defect is inefficient escape from LL; Delta and Notch trajectories enter organized *A*-field states more readily but show weaker HH self-persistence and greater leakage back to LL than wild type. The framework therefore helps separate pathway-level collapse, entry/progression into organized proneural dynamics, and maintenance of organized proneural-output states.

This has an important consequence for interpreting multicomponent nonlinear systems. DNOS and DPC are not only descriptors of final image complexity; when computed through time and across fields, they define a low-dimensional dynamical coordinate system for realized progression. In the developmental setting, genotypes can differ by endpoint, by trajectory length, by distance from wild-type trajectories, by the probability of leaving low-organization states, and by the conditional stability of high-organization states once reached. This is especially useful for developmental models in which visually similar final states may arise through different routes, or in which a mutant preserves one output field while disrupting an upstream signaling channel. In the proneural-wave model, this distinction is biologically central: a final *A*-field with non-zero complexity is not necessarily evidence of normal wave progression unless the trajectory and upstream multichannel state are also considered.

These observations indicate that DNOS and DPC can help summarize how reaction–diffusion and regulatory systems respond to changes in parameters, network structure, or perturbations. They do not replace geometry-specific analyses or mechanistic models. The same information could in principle be obtained from bespoke descriptors such as wavefront velocity, peak position, field-specific amplitude, autocorrelation length, or pathway-specific activity profiles. The value of the present framework is that it provides a common operational coordinate system in which endpoint state, field-specific organization, multichannel state organization, and dynamical route can be compared across perturbations and across distinct model classes without redesigning the descriptor for each case. The present paper deliberately keeps this comparison tied to the realized model fields themselves: endpoint, field-resolved, multichannel, and time-resolved analyses are all computed from the simulated variables under a fixed measurement protocol.

Future extensions might include applications to additional multicomponent fields, three-dimensional domains, experimental imaging data, and perturbation screens, as well as closer connections to bifurcation analysis and data-driven inference. More broadly, the framework developed here is intended as a tool for comparing realized organization in nonlinear pattern-forming systems: it provides a common language for assessing how different reaction–diffusion and regulatory systems realize spatial structure, how those outcomes change under parameter variation or perturbation, and how nonlinear patterning extends beyond what can be inferred from linear onset analysis alone. Within the present scope, DNOS and DPC define the core comparative space for this task, with UME serving as a supporting descriptor of uniformity where finer interpretive separation is required.

## 4 Methods

### 4.1 Numerical simulation and stability checks

All pattern-forming systems in this study were simulated on rectangular spatial grids using a standard five-point finite-difference discretization of the Laplacian and explicit time stepping. Unless stated otherwise for a specific model, we imposed Neumann (zero-flux) boundary conditions. Reaction terms were evaluated pointwise at each grid cell. For concentration-like variables, any small negative values induced numerically were truncated to 0.

For each parameter set, fields were initialized at or near the relevant homogeneous reference state with small-amplitude perturbations, and simulations were integrated to the model-specific final analysis time. Unless explicitly stated otherwise, DNOS, DPC, and UME were computed from the final snapshot only. The Notch–Delta–EGF analysis additionally used saved temporal checkpoints to construct time-resolved DNOS–DPC trajectories; no temporal interpolation was applied.

To verify that the reported patterns and complexity landscapes were not numerical artifacts, we repeated representative simulations with Δ*x* and Δ*t* halved while keeping the domain size and final time fixed. Spatial intensity statistics and compression-based complexity changed by less than 1%, and the locations of regime boundaries in the (DPC, DNOS, UME) feature space were unchanged. We additionally checked the discrete diffusion operator against the analytical spectrum of the Laplacian on test fields.

The linearized reaction–diffusion benchmark is used in two distinct ways. First, in Section 2.1, it is analyzed through its dispersion relation to obtain a spectral benchmark for homogeneous stability, finite-wavenumber Turing instability, growth rates, and dominant linear wavelengths. Second, in Section 2.3, the same linearized PDE is numerically integrated in time, and DNOS, DPC, and UME are computed from the resulting simulated pattern fields. For the Gray–Scott, FitzHugh–Nagumo, toggle-switch, and Notch–Delta–EGF systems, these measures are computed from simulated nonlinear pattern fields.

### 4.2 Spectral analysis of the linearized benchmark

To construct the linear benchmark shown in Fig. 1, we analyze the stability of a generic two-species reaction–diffusion system,

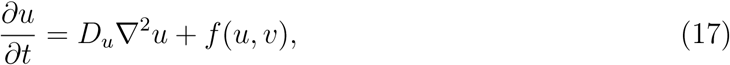

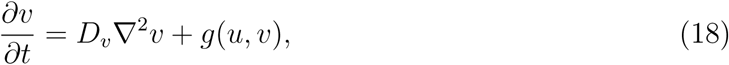

about a homogeneous steady state (*h, k*) satisfying

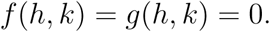

Linearizing around (*h, k*) gives

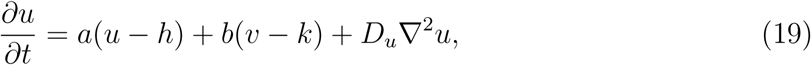

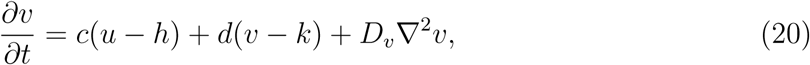

where

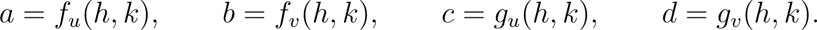

This spectral analysis is used only to characterize the onset of instability and the corresponding linear wavelength selection. It is therefore distinct from the time-evolution analysis in Section 2.3, where the same linearized PDE is numerically integrated and DNOS, DPC, and UME are computed from the resulting simulated pattern fields.

We consider perturbations of the form

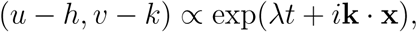

which lead to the *k*-dependent Jacobian

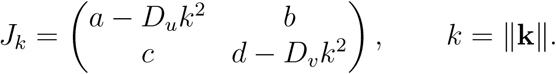

For each parameter set and wavenumber *k*, we compute the two eigenvalues *λ*_1_(*k*) and *λ*_2_(*k*) numerically.

The homogeneous steady state is linearly stable if

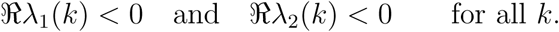

A diffusion-driven (Turing) instability occurs when the state is stable at *k* = 0 but unstable for some finite *k >* 0. For a standard two-species system with ordinary diffusion, this finite-wavenumber instability is the relevant linear signature of Turing pattern formation. Accordingly, we restrict the analysis to quantities that are directly supported by the dispersion relation, rather than assigning nonlinear bifurcation labels such as pitchfork or transcritical classes.

The diffusion coefficients *D_u_* and *D_v_* were varied on a logarithmic grid in the range 10^−5^ to 10^−3^, while the Jacobian entries *a, b, c, d* were sampled uniformly from [−2, 2]. For each parameter set, we evaluated the dispersion relation on a grid of wavenumbers *k* ∈ [0, 100]. For each diffusion pair (*D_u_, D_v_*), we computed:

1. the fraction of sampled Jacobians stable at *k* = 0;
2. the fraction that were Turing-unstable in the numerical dispersion scan, i.e. stable at *k* = 0 but unstable for some finite *k >* 0;
3. the mean maximum growth rate max max

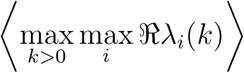

over the Turing-unstable cases;
4. the mean dominant wavelength

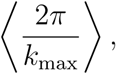

where *k*_max_ is the wavenumber that maximises the growth rate for a given unstable sample.

We also evaluated the classical analytic Turing conditions as a validation of the numerical dispersion scan. These validation results are not used as the main figure in the manuscript but are reported in Supplementary Fig. S1 of the supplementary material.

This spectral benchmark identifies where diffusion-driven instabilities are expected, how strongly unstable modes grow, and which linear wavelengths are selected at onset. It does not, however, determine the realized long-time pattern morphology. For that reason, the main text complements the dispersion-relation analysis with direct complexity measurements on simulated pattern fields in Section 2.3 and in the nonlinear model classes analyzed subsequently.

### 4.3 Complexity measures

We quantify realized patterns using three complementary measures: the Diversity of Number of States (DNOS), the Diversity of Pattern Complexity (DPC), and the Uniformity Measure of Entropy (UME). All three operate directly on simulated spatial fields after normalization to the unit interval and do not require geometric feature extraction.

For selected robustness analyses reported in the supplementary material, parameter sets were stratified into five non-overlapping pools and summary statistics were computed across those pools. The main results in the paper, however, are based on per-pattern measures and on the regime structure they induce across parameter space.

Table 5 summarizes the fixed measurement protocol used for the main analyses. The same protocol is used across the model classes unless a figure caption or model-specific Methods subsection states otherwise.

**Table 5:**
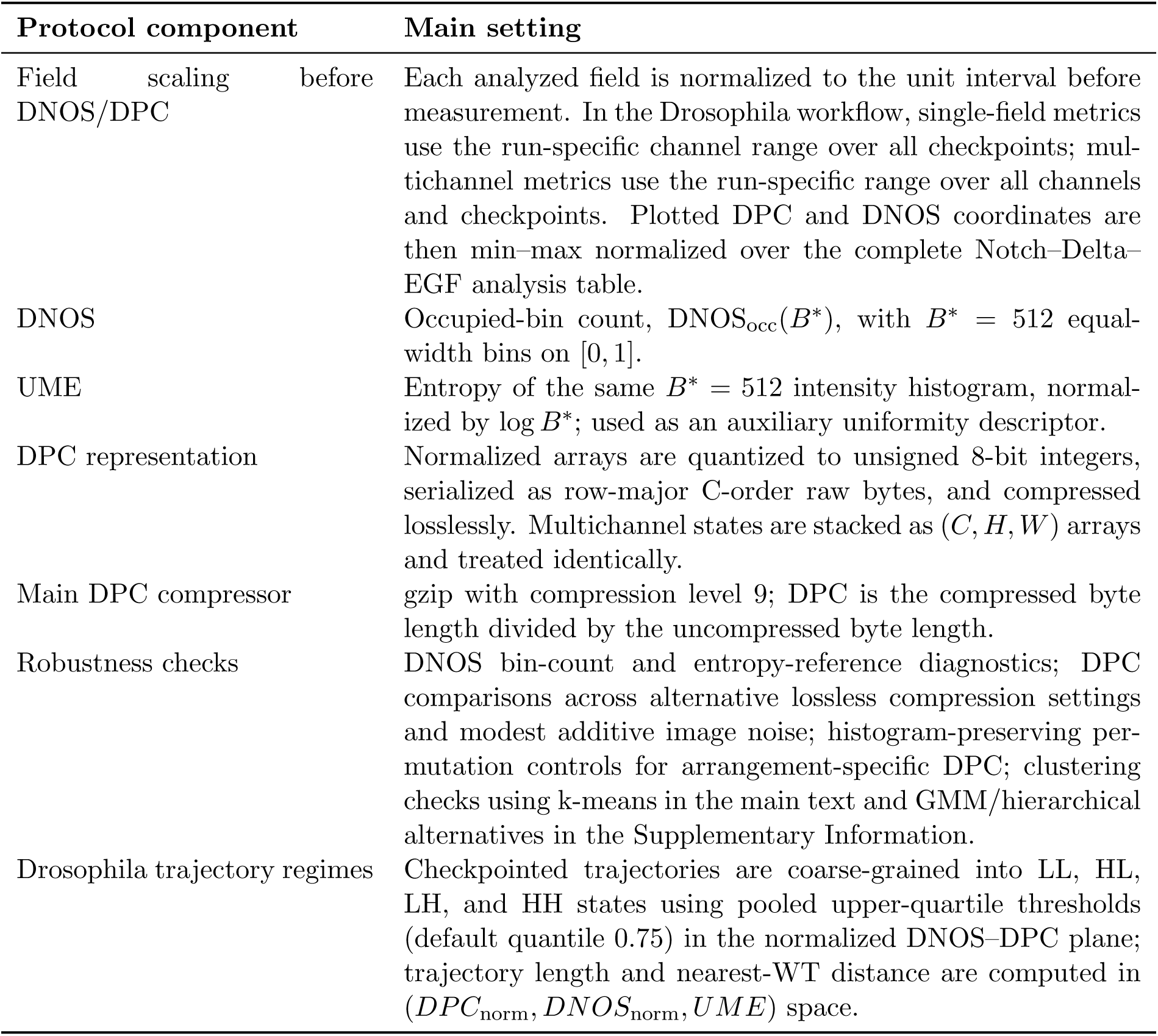
Measurement protocol used for the main DNOS–DPC analyses.

### 4.4 Diversity of Number of States (DNOS)

Let *X*(**x**) ∈ [0, 1] denote a normalized concentration field on a regular grid. We partition the interval [0, 1] into *B* equal-width bins and compute the empirical histogram with probabilities *p_i_*. Our operational DNOS is the number of occupied bins,

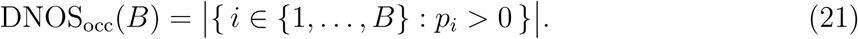

This quantity measures how broadly a pattern explores its available intensity space at the chosen resolution.

Throughout the main analyses we fix a high-resolution bin count *B*^∗^ = 512 and use DNOS_occ_(*B*^∗^) as the DNOS coordinate in feature space. Patterns concentrated into a narrow intensity range occupy fewer bins and therefore have lower DNOS, whereas patterns that spread across a broader range of intensities occupy more bins and therefore have higher DNOS. Entropy-based and bin-free DNOS diagnostics were used only to verify the consistency of this occupied-bin definition and are reported in the Supplementary Information.

### 4.5 Diversity of Pattern Complexity (DPC)

To quantify structural complexity, we estimate the algorithmic compressibility of each pattern using lossless compression. In the main implementation, a normalized field *X* ∈ [0, 1] is quantized to unsigned 8-bit integers as *Q* = round(255*X*), serialized as a row-major C-order byte string, and compressed with gzip at compression level 9. For the resulting byte representation *I*, DPC is

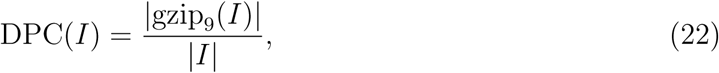

where |*I*| is the number of uncompressed bytes and |gzip_9_(*I*)| is the number of compressed bytes. Multichannel states are stacked as arrays of shape (*C, H, W*) before the same normalization, quantization, serialization, and compression procedure is applied. Low DPC values correspond to highly compressible, regular, or nearly homogeneous patterns, whereas high DPC values indicate less compressible and therefore structurally richer patterns.

DPC is computed independently for each pattern and used directly as a per-pattern scalar quantity throughout the paper. The gzip/uint8/raw-byte setting is the main measurement protocol. Robustness checks using alternative lossless compression settings, alternative discretizations, and modest image noise confirmed that the ordering induced by DPC is stable; these diagnostics are reported in the Supplementary Information.

### 4.6 Uniformity Measure of Entropy (UME)

UME quantifies how evenly a pattern uses its available intensity space. Given the same binned intensity distribution used for DNOS, we define

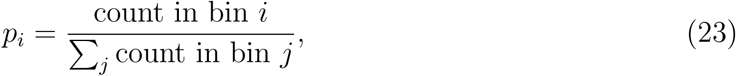

count in bin *j*

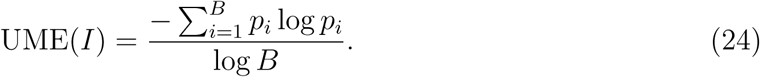

UME equals 1 for a perfectly uniform histogram and 0 when all mass is concentrated in a single bin.

In practice, UME is computed using the same high-resolution binning *B*^∗^ as DNOS. DNOS therefore records how many bins are occupied, whereas UME records how evenly those occupied bins are filled. When UME is applied to binned spatial intensities we denote it by UME(*S*), and when it is applied to binned per-pattern compression ratios we denote it by UME(CR).

In the main analyses, UME is used as an auxiliary descriptor to refine separation between regimes with similar DNOS but different degrees of uniformity. In particular, UME(*S*) summarizes how evenly intensity space is occupied, while UME(CR) provides the corresponding uniformity summary in complexity space. Their joint use in the cross-model comparison is described in Section 2.7.

### 4.7 Field-resolved, multichannel, and time-resolved DNOS–DPC analysis

The Notch–Delta–EGF analysis begins with the same static endpoint DNOS–DPC framework used elsewhere in the paper, preserving the final-snapshot genotype comparison as a baseline. It then extends the endpoint framework in three ways. First, measures are computed separately for each biological field *E*, *N*, *D*, and *A*, allowing pathway-level and output-level organization to be distinguished. Second, the four fields are analyzed as a multichannel object [*E, N, D, A*], which represents the full signaling state of the tissue. Third, measures are computed at saved temporal checkpoints, converting each run into a trajectory through complexity space. The released code implements this workflow as a scripted pipeline: checkpointed simulations are stored in a tensor with axes run, checkpoint, channel, row, and column; the analysis stage writes a tidy per-checkpoint metric table; and the plotting stage reads only these tables to generate endpoint, trajectory, and transition figures.

For a field *X_r_*(*t*) from run *r* at checkpoint *t*, we compute

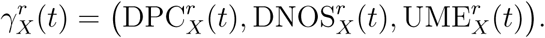

Endpoint figures use 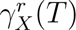 at the final checkpoint *T*. Trajectory figures use the full ordered sequence 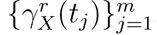. DPC and DNOS are min–max normalized over the complete Notch–Delta–EGF analysis before plotting, so that field-specific and multichannel summaries share a common visual scale.

#### Algorithm 1 Field-resolved and multichannel endpoint DNOS-DPC landscape

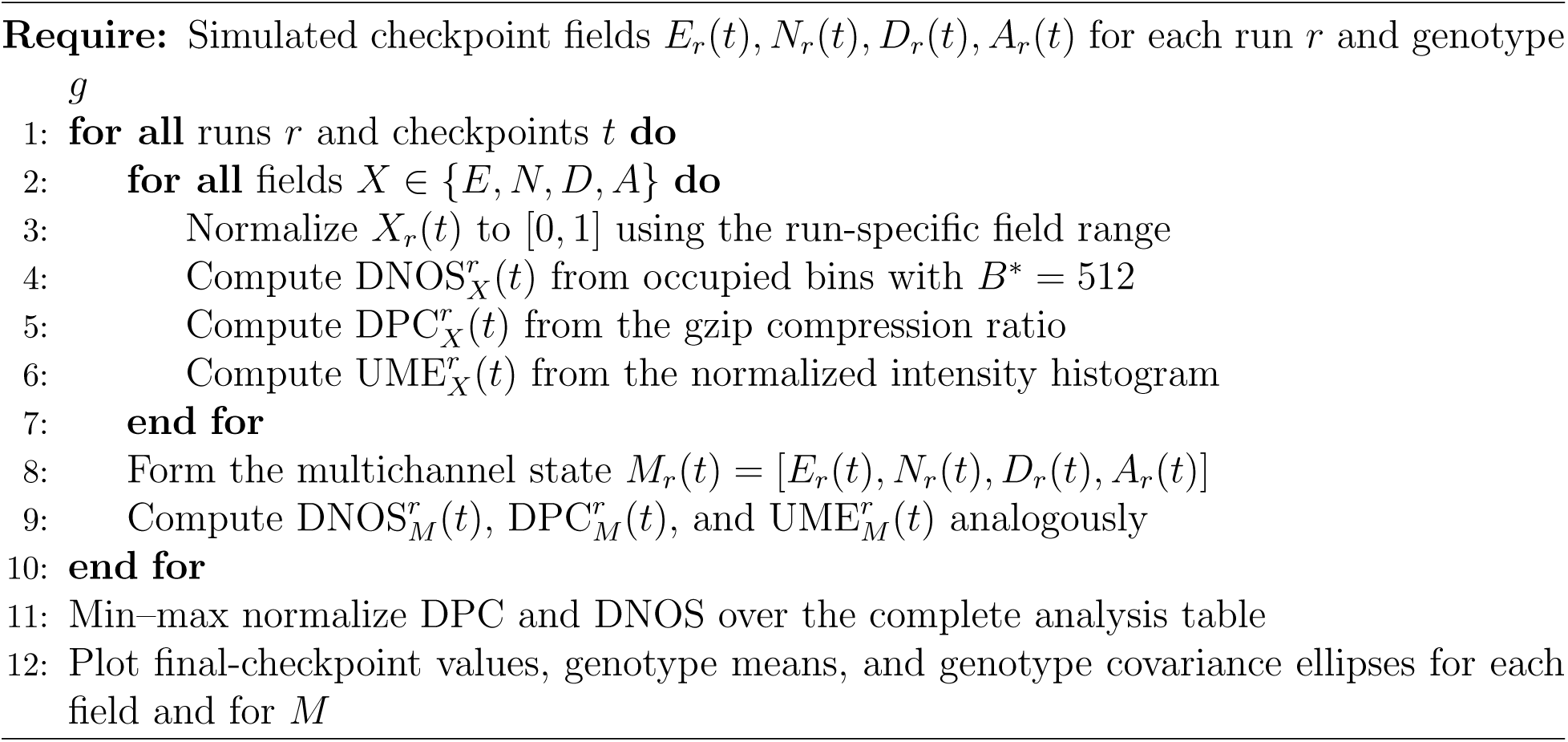

To distinguish structural arrangement from the intensity histogram alone, we also computed an arrangement-conditioned DPC diagnostic for selected endpoint analyses. For each pattern, pixel values were randomly permuted to generate histogram-preserving surrogate images; in the released analysis workflow, the default is 32 final-checkpoint surrogates per analyzed pattern. The arrangement-specific score was then computed as

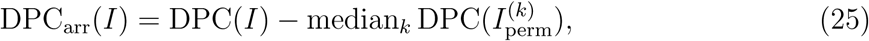

with the corresponding ratio and *z*-score used as robustness diagnostics. This diagnostic is used to verify that high DPC is not driven only by broad intensity occupancy, but also by spatial arrangement.

#### Algorithm 2 Histogram-conditioned arrangement-specific DPC diagnostic

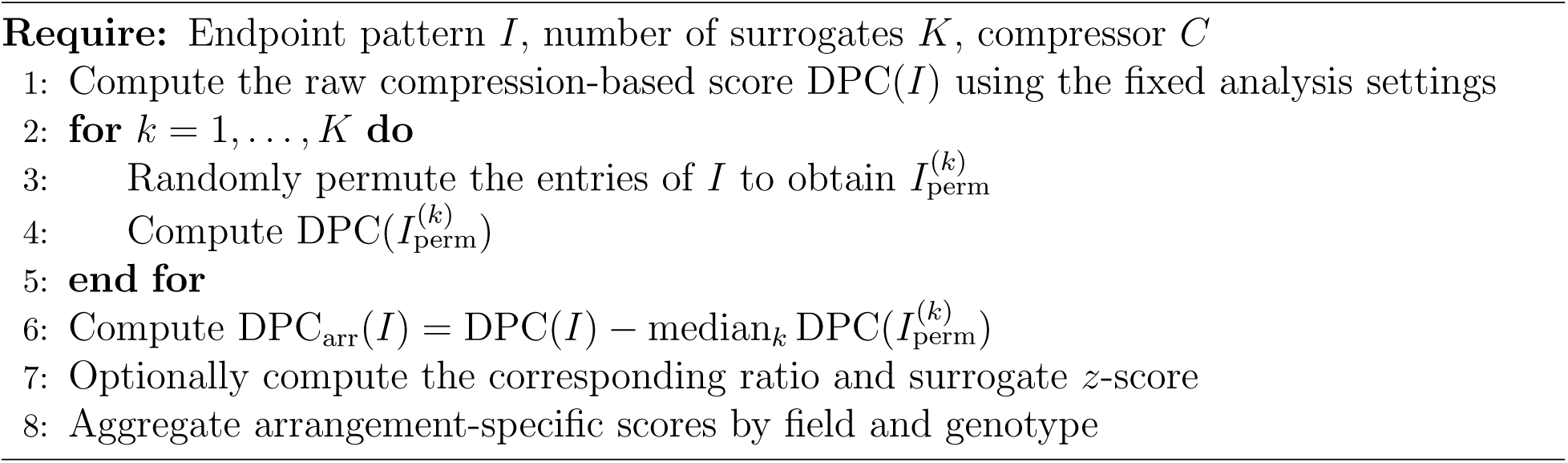

#### Algorithm 3 Time-resolved trajectories and coarse-grained regime transitions

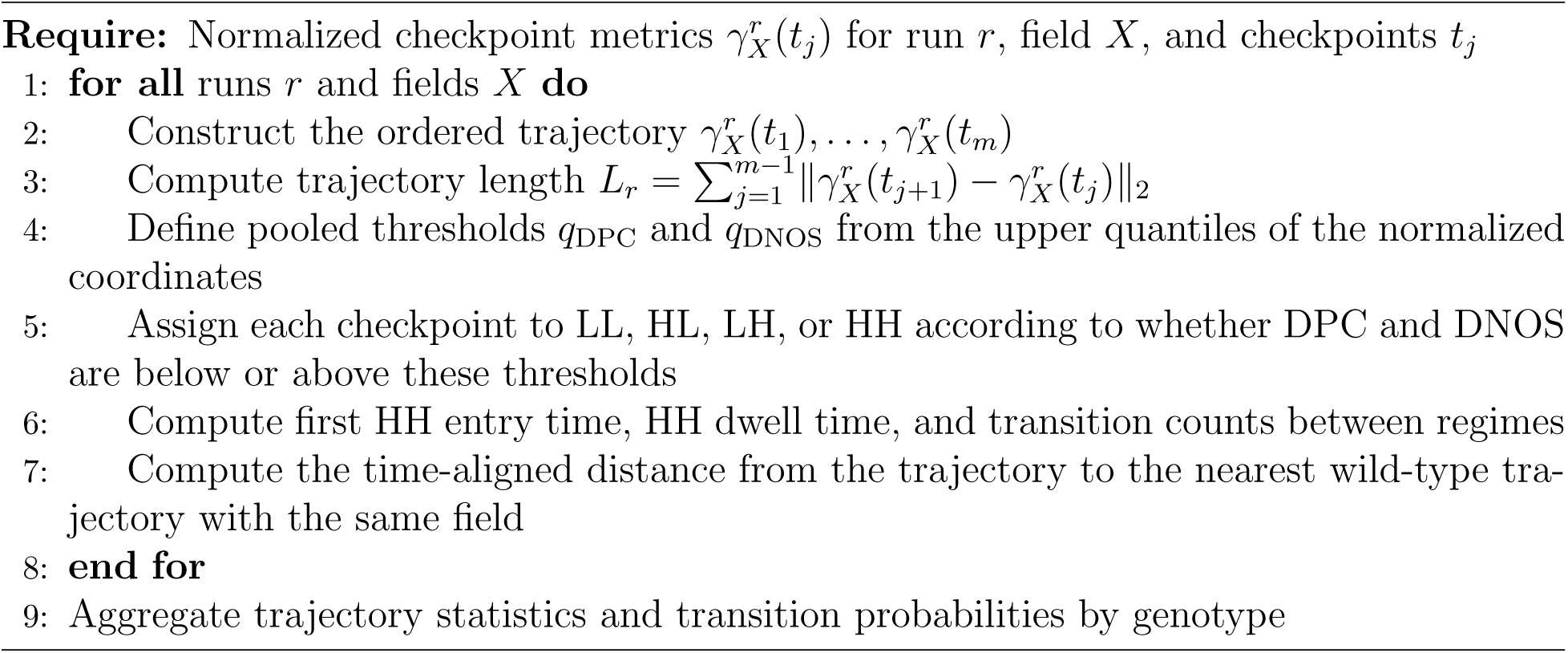

### 4.8 Regime identification

Regimes were identified by clustering patterns in the feature spaces used in the corresponding figures. Throughout the main paper, all displayed clustering results use *k*-means, consistent across the linearized reaction–diffusion, Gray–Scott, FitzHugh–Nagumo, and toggle-switch analyses. Depending on the figure, clustering was performed either in the (DPC, DNOS) plane or in the augmented (DPC, DNOS, UME) space after *z*-scaling each component.

The number of clusters was chosen separately for each analysis and is reported in the corresponding Results subsection and figure caption. The purpose of clustering is not to recover visually defined pattern classes, but to identify stable computational regimes in the low-dimensional feature space induced primarily by DNOS and DPC, with UME included only where explicitly stated as a supporting descriptor. Alternative clustering methods, including Gaussian mixture models and hierarchical clustering, yielded essentially equivalent partitions and are reported only as robustness checks in the Supplementary Information.

### 4.9 Auxiliary measures for benchmarking DPC

Expanded definitions of the auxiliary benchmarking measures RDH, ESP, and WLE are provided in Sec. S6 of the supplementary material.

### 4.10 Finite-difference implementation

For a generic two-species system

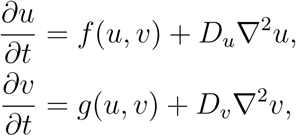

we discretize space on a regular grid with Δ*x* = Δ*y* and approximate the Laplacian by the standard five-point stencil,

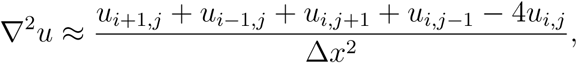

and analogously for *v*. Time stepping uses forward Euler updates,

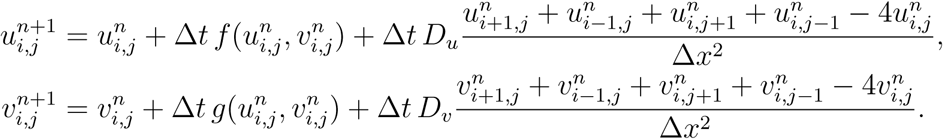

This scheme is used for both the linearized benchmark in Section 2.3 and the nonlinear reaction–diffusion models analyzed elsewhere in the paper; for multicomponent systems such as the Notch–Delta–EGF model, the same update is applied componentwise. Because the scheme is explicit, Δ*t* was chosen sufficiently small relative to Δ*x* and to the relevant diffusion scale. The truncation error is *O*(Δ*x*^2^) in space and *O*(Δ*t*) in time.

## Supplementary Material

See the supplementary material for additional details on the linear spectral benchmark, DNOS diagnostics, DPC robustness checks, UME, clustering robustness, and auxiliary bench-marking measures.

## Author Declarations

### Ethics Approval

Not applicable. This study used computational simulations and did not involve experiments with human participants or animals.

### Author Contributions

Jurgen Riedel: Conceptualization; Methodology; Software; Formal analysis; Investigation; Data curation; Visualization; Writing – original draft. Chris P. Barnes: Supervision; Conceptualization; Writing – review and editing. Both authors reviewed and approved the final manuscript.

### Data Availability

The data that support the findings of this study are openly available in Zenodo at https://doi.org/10.5281/zenodo.14525543.

## References

[1] Alan M. Turing. The chemical basis of morphogenesis. Philosophical Transactions of the Royal Society of London. Series B, Biological Sciences, 237(641):37–72, 1952. doi: 10.1098/rstb.1952.0012. URL https://doi.org/10.1098/rstb.1952.0012.

[2] Alfred Gierer and Hans Meinhardt. A theory of biological pattern formation. Kybernetik, 12(1):30–39, 1972. doi: 10.1007/BF00289234. URL https://doi.org/10.1007/BF00289234.

[3] Michael F. Staddon. How the zebra got its stripes: Curvature-dependent diffusion orients turing patterns on three-dimensional surfaces. Physical Review E, 110(3):034402, 2024. doi: 10.1103/PhysRevE.110.034402.

[4] Shigeru Kondo, Masakatsu Watanabe, and Seita Miyazawa. Studies of turing pattern formation in zebrafish skin. Philosophical Transactions of the Royal Society A: Mathematical, Physical and Engineering Sciences, 379(2216):20200274, 2021. doi: 10.1098/rsta.2020.0274. URL https://doi.org/10.1098/rsta.2020.0274.

[5] Akiko Nakamasu, Go Takahashi, Akio Kanbe, and Shigeru Kondo. Interactions between zebrafish pigment cells responsible for the generation of turing patterns. Proceedings of the National Academy of Sciences, 106(21):8429–8434, 2009. doi: 10.1073/pnas.0808622106.

[6] Ebrahim Jahanbakhsh and Michel C. Milinkovitch. Modeling convergent scale-by-scale skin color patterning in multiple species of lizards. Current Biology, 32(23):R1306–R1308, 2022. doi: 10.1016/j.cub.2022.10.044.

[7] Philip K. Maini and Thomas E. Woolley. The Turing model for biological pattern formation. In Alessandra Bianchi, Thomas Hillen, Mark Lewis, and Yingfei Yi, editors, The Dynamics of Biological Systems, volume 4 of Mathematics of Planet Earth. Springer, Cham, 2019. doi: 10.1007/978-3-030-22583-47. URL https://doi.org/10.1007/978-3-030-22583-4_7.

[8] Benjamin M. Alessio and Ankur Gupta. Diffusiophoresis-enhanced Turing patterns. Science Advances, 9(45):eadj2457, 2023. doi: 10.1126/sciadv.adj2457. URL https://doi.org/10.1126/sciadv.adj2457.

[9] Philip K. Maini. Using mathematical models to help understand biological pattern formation. Comptes Rendus Biologies, 327(3):225–234, 2004. doi: 10.1016/j.crvi.2003.05.006. URL https://doi.org/10.1016/j.crvi.2003.05.006.

[10] Vaclav Klika. Significance of non-normality-induced patterns: Transient growth versus asymptotic stability. Chaos: An Interdisciplinary Journal of Nonlinear Science, 27(7): 073120, 2017. doi: 10.1063/1.4985256. URL https://doi.org/10.1063/1.4985256.

[11] Sean T. Vittadello, Thomas Leyshon, David Schnoerr, and Michael P. H. Stumpf. Turing pattern design principles and their robustness. Philosophical Transactions of the Royal Society A: Mathematical, Physical and Engineering Sciences, 379(2213):20200272, 2021. doi: 10.1098/rsta.2020.0272. URL https://doi.org/10.1098/rsta.2020.0272.

[12] Andrew L. Krause, Eamonn A. Gaffney, Thomas J. Jewell, Vaclav Klika, and Benjamin J. Walker. Turing instabilities are not enough to ensure pattern formation. Bulletin of Mathematical Biology, 86:21, 2024. doi: 10.1007/s11538-023-01250-4. URL https://doi.org/10.1007/s11538-023-01250-4.

[13] Yanyan Chen and Javier Buceta. A non-linear analysis of turing pattern formation. PLoS ONE, 14(8):e0220994, 08 2019. doi: 10.1371/journal.pone.0220994. URL https://doi.org/10.1371/journal.pone.0220994.

[14] J. L. Aragon, R. A. Barrio, T. E. Woolley, R. E. Baker, and P. K. Maini. Nonlinear effects on Turing patterns: Time oscillations and chaos. Physical Review E, 86(2): 026201, 2012. doi: 10.1103/PhysRevE.86.026201. URL https://doi.org/10.1103/PhysRevE.86.026201.

[15] Martina Oliver Huidobro and Robert G. Endres. Effects of multistability, absorbing boundaries and growth on turing pattern formation. bioRxiv, 2024. doi: 10.1101/2024.09.09.611947. URL https://doi.org/10.1101/2024.09.09.611947.

[16] Roozbeh H. Pazuki and Robert G. Endres. Robustness of turing models and gene regulatory networks with a sweet spot. Phys. Rev. E, 109:064305, 2024. doi: 10.1103/PhysRevE.109.064305. URL https://doi.org/10.1103/PhysRevE.109.064305.

[17] Shibashis Paul, Joy Adetunji, and Tian Hong. Widespread biochemical reaction networks enable turing patterns without imposed feedback. Nature Communications, 15:8380, 2024. doi: 10.1038/s41467-024-52591-0. URL https://doi.org/10.1038/s41467-024-52591-0.

[18] Jasper van der Kolk, Guillermo Garcıa-Perez, Nikos E. Kouvaris, M. Angeles Serrano, and Marian Boguna. Emergence of geometric Turing patterns in complex networks. Physical Review X, 13(2):021038, 2023. doi: 10.1103/PhysRevX.13.021038. URL https://doi.org/10.1103/PhysRevX.13.021038.

[19] Ajay Subbaroyan, Olivier C. Martin, and Areejit Samal. Minimum complexity drives regulatory logic in boolean models of living systems. PNAS Nexus, 1(1):pgac017, 2022. doi: 10.1093/pnasnexus/pgac017.

[20] Pascal F. Hagolani, Roland Zimm, Renske Vroomans, and Isaac Salazar-Ciudad. On the evolution and development of morphological complexity: A view from gene regulatory networks. PLOS Computational Biology, 17(2):e1008570, 2021. doi: 10.1371/journal.pcbi.1008570. URL https://doi.org/10.1371/journal.pcbi.1008570.

[21] Saverio Forestiero. The historical nature of biological complexity and the ineffectiveness of the mathematical approach to it. Acta Biotheoretica, 2022. doi: 10.1007/s12064-022-00369-7. URL https://doi.org/10.1007/s12064-022-00369-7.

[22] S. Miyazawa, M. Okamoto, and S. Kondo. Blending of animal colour patterns by hybridization. Nature Communications, 1(66):1–7, 2010. doi: 10.1038/ncomms1071.

[23] D.E. Lee, D.R. Cavener, and M.L. Bond. Seeing spots: quantifying mother-offspring similarity and assessing fitness consequences of coat pattern traits in a wild population of giraffes. PeerJ, 6:e5690, 2018. doi: 10.7717/peerj.5690.

[24] T. Glimm, M. Kiskowski, N. Moreno, and Y. Chiari. Capturing and analyzing pattern diversity: an example using the melanistic spotted patterns of leopard geckos. PeerJ, 9:e11829, 2021. doi: 10.7717/peerj.11829.

[25] G.H. Moro, M. de Gomensoro Malheiros, and M. Walter. Quantitative descriptors for a range of visual biologic pigmentation patterns. In 35th SIBGRAPI Conference on Graphics, Patterns and Images, pages 133–138, 2022. doi: 10.1109/SIBGRAPI55357.2022.9991769.

[26] B. Ermentrout. Stripes or spots? nonlinear effects in bifurcation of reaction–diffusion equations on the square. Proceedings of the Royal Society of London. Series A: Mathematical and Physical Sciences, 434(1891):413–417, 1991. doi: 10.1098/rspa.1991.0100. URL https://doi.org/10.1098/rspa.1991.0100.

[27] T.J. Callahan and E. Knoblauch. Pattern formation in reaction–diffusion systems: a bifurcation analysis. Journal of Mathematical Biology, 48(4):385–417, 2004. doi: 10.1007/s00285-003-0240-8.

[28] P. Gray and S. K. Scott. Autocatalytic reactions in the isothermal, continuous stirred tank reactor: isolas and other forms of multistability. Chemical Engineering Science, 38(1):29–43, 1983. doi: 10.1016/0009-2509(83)80194-7.

[29] John E. Pearson. Complex patterns in a simple system. Science, 261(5118):189–192, 1993. doi: 10.1126/science.261.5118.189.

[30] Robert E. Munafo. Generalized Gray–Scott models. Complex Systems, 27(4):315–367, 2018. doi: 10.25088/ComplexSystems.27.4.315.

[31] John J. Mueller and Eshel Ben-Jacob. Dissipative structures and morphogenesis in bacterial colonies. Physica A, 249(1–4):123–130, 1998. doi: 10.1016/S0378-4371(97)00568-3.

[32] J. S. McGough and K. Riley. Pattern formation in the gray–scott model. Nonlinear Analysis: Real World Applications, 5(1):105–121, 2004. doi: 10.1016/S1468-1218(03)00020-8.

[33] Satyvir Singh, R. C. Mittal, Shafeeq Rahman Thottoli, and Mohammad Tamsir. High-fidelity simulations for Turing pattern formation in multi-dimensional Gray–Scott reaction-diffusion system. Applied Mathematics and Computation, 452:128079, 2023. doi: 10.1016/j.amc.2023.128079. URL https://doi.org/10.1016/j.amc.2023.128079.

[34] Wenjing Jiang, Ziling Lu, and Jian Wang. Uniform patterns formation based on Gray–Scott model for 3D printing. Computer Physics Communications, 295:108974, 2024. doi: 10.1016/j.cpc.2023.108974. URL https://doi.org/10.1016/j.cpc.2023.108974.

[35] Alan L. Hodgkin and Andrew F. Huxley. A quantitative description of membrane current and its application to conduction and excitation in nerve. The Journal of Physiology, 117(4):500–544, 1952. doi: 10.1113/jphysiol.1952.sp004764. URL https://doi.org/10.1113/jphysiol.1952.sp004764.

[36] Richard FitzHugh. Impulses and physiological states in theoretical models of nerve membrane. Biophysical Journal, 1(6):445–466, 1961. doi: 10.1016/S0006-3495(61)86902-6. URL https://doi.org/10.1016/S0006-3495(61)86902-6.

[37] Jinichi Nagumo, Suguru Arimoto, and Shuji Yoshizawa. An active pulse transmission line simulating nerve axon. Proceedings of the IRE, 50(10):2061–2070, 1962. doi: 10.1109/JRPROC.1962.288235. URL https://doi.org/10.1109/JRPROC.1962.288235.

[38] Gaetana Gambino, Maria Cristina Lombardo, Gino Rubino, and Marcello Sammartino. Pattern selection in the 2d fitzhugh–nagumo model. Ricerche di Matematica, 67(2): 377–392, 2018. doi: 10.1007/s11587-018-0424-6.

[39] Daniel Cebrian-Lacasa, Pedro Parra-Rivas, Daniel Ruiz-Reynes, and Lendert Gelens. Six decades of the FitzHugh–Nagumo model: A guide through its spatiotemporal dynamics and influence across disciplines. Physics Reports, 1096:1–39, 2024. doi: 10.1016/j.physrep.2024.09.014. URL https://doi.org/10.1016/j.physrep.2024.09.014.

[40] Ben D. MacArthur, Avi Ma’ayan, and Ihor R. Lemischka. Systems biology of stem cell fate and cellular reprogramming. Nature Reviews Molecular Cell Biology, 10(9): 672–681, 2009. doi: 10.1038/nrm2766. URL https://doi.org/10.1038/nrm2766.

[41] Johan Paulsson. Summing up the noise in gene networks. Nature, 427(6973):415–418, 2004. doi: 10.1038/nature02257. URL https://doi.org/10.1038/nature02257.

[42] Ingo Roeder and Ingmar Glauche. Towards an understanding of lineage specification in hematopoietic stem cells: A mathematical model for the interaction of transcription factors GATA-1 and PU.1. Journal of Theoretical Biology, 241(4):852–865, 2006. doi: 10.1016/j.jtbi.2006.01.021. URL https://doi.org/10.1016/j.jtbi.2006.01.021.

[43] J. Lynn Cherry and Frederick R. Adler. How to make a biological switch. Journal of Theoretical Biology, 203(2):117–133, 2000. doi: 10.1006/jtbi.2000.1068. URL https://doi.org/10.1006/jtbi.2000.1068.

[44] Campbell Duff, Kate Smith-Miles, Leo Lopes, and Tianhai Tian. Mathematical modelling of stem cell differentiation: The PU.1–GATA-1 interaction. Journal of Mathematical Biology, 64(3):449–468, 2012. doi: 10.1007/s00285-011-0419-3. URL https://doi.org/10.1007/s00285-011-0419-3.

[45] Ruben Perez-Carrasco, Pilar Guerrero, James Briscoe, and Karen Page. Intrinsic noise profoundly alters the dynamics and steady state of morphogen-controlled bistable genetic switches. PLOS Computational Biology, 12(10):e1005154, 2016. doi: 10.1371/journal.pcbi.1005154. URL https://doi.org/10.1371/journal.pcbi.1005154.

[46] Patrick B. Warren and Pieter Rein ten Wolde. Statistical analysis of the spatial patterns of cell–cell communication. Journal of Theoretical Biology, 231(4):563–579, 2004. doi: 10.1016/j.jtbi.2004.07.012. URL https://doi.org/10.1016/j.jtbi.2004.07.012.

[47] Thomas B. Kepler and Timothy C. Elston. Stochasticity in transcriptional regulation: Origins, consequences, and mathematical representations. Biophysical Journal, 81(6): 3116–3136, 2001. doi: 10.1016/S0006-3495(01)75949-8. URL https://doi.org/10.1016/S0006-3495(01)75949-8.

[48] Vijay Chickarmane, Tariq Enver, and Carsten Peterson. Computational modeling of the hematopoietic erythroid–myeloid switch reveals insights into cooperativity, priming, and irreversibility. PLOS Computational Biology, 5(1):e1000268, 2009. doi: 10.1371/journal.pcbi.1000268. URL https://doi.org/10.1371/journal.pcbi.1000268.

[49] Michael K. Strasser, Fabian J. Theis, and Carsten Marr. Stability and multiattractor dynamics of a toggle switch based on a two-stage model of stochastic gene expression. Biophysical Journal, 102(1):19–29, 2012. doi: 10.1016/j.bpj.2011.11.4000. URL https://doi.org/10.1016/j.bpj.2011.11.4000.

[50] Ming-Chang Huang, Jinn-Wen Wu, Yu-Pin Luo, and Karen G. Petrosyan. Fluctuations in gene regulatory networks as Gaussian colored noise. The Journal of Chemical Physics, 132(15):155101, 2010. doi: 10.1063/1.3385468. URL https://doi.org/10.1063/1.3385468.

[51] Ushasi Roy, Divyoj Singh, Navin Vincent, Chinmay K. Haritas, and Mohit Kumar Jolly. Spatiotemporal patterning enabled by gene regulatory networks. ACS Omega, 8(4):3713–3725, 2023. doi: 10.1021/acsomega.2c04581. URL https://doi.org/10.1021/acsomega.2c04581.

[52] Makoto Sato, Tetsuo Yasugi, Yoshiaki Minami, and Masaharu Nagayama. Notch-mediated lateral inhibition regulates proneural wave propagation when combined with egf-mediated reaction diffusion. Proceedings of the National Academy of Sciences, 113(35):E5153–E5162, August 2016. doi: 10.1073/pnas.1602739113. URL https://doi.org/10.1073/pnas.1602739113.

[53] Makoto Sato and Tetsuo Yasugi. Regulation of proneural wave propagation through a combination of notch-mediated lateral inhibition and egf-mediated reaction diffusion. In Current Topics in Developmental Biology / Advances in Experimental Medicine and Biology, volume 1218 of Advances in Experimental Medicine and Biology, pages 77–91. 2020. doi: 10.1007/978-3-030-34436-85. URL https://doi.org/10.1007/978-3-030-34436-8_5.

[54] David J. Jorg, Elizabeth E. Caygill, Anna E. Hakes, Esteban G. Contreras, Andrea H. Brand, and Benjamin D. Simons. The proneural wave in the drosophila optic lobe is driven by an excitable reaction-diffusion mechanism. eLife, 8:e40919, 2019. doi: 10.7554/eLife.40919. URL https://doi.org/10.7554/eLife.40919.

[55] Miaoxing Wang, Xujun Han, Chuyan Liu, Rie Takayama, Tetsuo Yasugi, Shin-Ichiro Ei, Masaharu Nagayama, Yoshitaro Tanaka, and Makoto Sato. Intracellular trafficking of notch orchestrates temporal dynamics of notch activity in the fly brain. Nature Communications, 12(1):2083, 2021. doi: 10.1038/s41467-021-22442-3. URL https://doi.org/10.1038/s41467-021-22442-3.

[56] Tetsuo Yasugi and Makoto Sato. Mathematical modeling of notch dynamics in drosophila neural development. Fly, 16(1):24–36, 2022. doi: 10.1080/19336934.2021.1953363. URL https://doi.org/10.1080/19336934.2021.1953363.

